# Epstein-Barr Virus Orchestrates Spatial Reorganization and Immunomodulation within the Classic Hodgkin Lymphoma Tumor Microenvironment

**DOI:** 10.1101/2024.03.05.583586

**Authors:** Yao Yu Yeo, Huaying Qiu, Yunhao Bai, Bokai Zhu, Yuzhou Chang, Jason Yeung, Hendrik A. Michel, Kyle Wright, Muhammad Shaban, Sam Sadigh, Dingani Nkosi, Vignesh Shanmugam, Philip Rock, Stephanie Pei Tung Yiu, Precious Cramer, Julia Paczkowska, Pierre Stephan, Guanrui Liao, Amy Y. Huang, Hongbo Wang, Han Chen, Leonie Frauenfeld, Bidisha Mitra, Benjamin E. Gewurz, Christian M. Schürch, Bo Zhao, Garry P. Nolan, Baochun Zhang, Alex K. Shalek, Michael Angelo, Faisal Mahmood, Qin Ma, W. Richard Burack, Margaret A. Shipp, Scott J. Rodig, Sizun Jiang

## Abstract

Classic Hodgkin Lymphoma (cHL) is a tumor composed of rare malignant Hodgkin and Reed-Sternberg (HRS) cells nested within a T-cell rich inflammatory immune infiltrate. cHL is associated with Epstein-Barr Virus (EBV) in 25% of cases. The specific contributions of EBV to the pathogenesis of cHL remain largely unknown, in part due to technical barriers in dissecting the tumor microenvironment (TME) in high detail. Herein, we applied multiplexed ion beam imaging (MIBI) spatial pro-teomics on 6 EBV-positive and 14 EBV-negative cHL samples. We identify key TME features that distinguish between EBV-positive and EBV-negative cHL, including the relative predominance of memory CD8 T cells and increased T-cell dysfunction as a function of spatial proximity to HRS cells. Building upon a larger multi-institutional cohort of 22 EBV-positive and 24 EBV-negative cHL samples, we orthogonally validated our findings through a spatial multi-omics approach, coupling whole transcriptome capture with antibody-defined cell types for tu-mor and T-cell populations within the cHL TME. We delineate contrasting transcriptomic immunological signatures between EBV-positive and EBV-negative cases that differently impact HRS cell proliferation, tumor-immune interactions, and mecha-nisms of T-cell dysregulation and dysfunction. Our multi-modal framework enabled a comprehensive dissection of EBV-linked reorganization and immune evasion within the cHL TME, and highlighted the need to elucidate the cellular and molecular fac-tors of virus-associated tumors, with potential for targeted therapeutic strategies.

## Introduction

Classic Hodgkin Lymphoma (cHL) is a B-cell lymphoid malignancy that most frequently occurs in young adults and less commonly in the elderly (1). The cHL tumor microenvironment (TME) has unique features, including rare malignant Hodgkin and Reed-Sternberg (HRS) multinucleated cells (comprising 1-5% of the cHL TME cell composition) nested within a dense T-cell rich inflammatory immune cell infiltrate. Although the etiology of cHL remains to be defined, cHL is associated with the oncogenic Epstein-Barr Virus (EBV) in approximately 25% of cases (2–4). Of note, cHL has lower cure rates and is more frequently EBV-positive in older patients (1). It remains elusive as to whether this is due to poorer tolerance of aggressive chemotherapy, or inherent biological differences associated with EBV infection.

EBV-positive and EBV-negative HRS cells are morphologically indistinguishable, but emerging evidence suggests that the cHL TME is modulated by the presence of the virus (5). EBV generally establishes latency II in HRS cells to express oncoproteins, including latent membrane protein 1 (LMP1), LMP2A, and EBV nuclear antigen 1 (EBNA1). LMP1 and EBNA1 expression in HRS cells is associated with the upregulation of immunomodulatory cytokines including CXCL10 (6), IL-6 (7), IL-10 (8, 9), and CCL20 (10), secreted factors which can significantly alter immune cell responses and organization within the TME (11). EBV-positive HRS cells also retain MHC Class I expression more frequently than their virus-negative counterparts, which often have a genetic basis for beta-2 microglobulin (B2M) loss and a higher mutational burden (12–15). These results are consistent with the ability of EBV-positive HRS cells that retain MHC Class I expression to present latent EBV oncoprotein antigens (16), and potential increase in CD8 T-cell activation within the EBV-positive cHL TME (17).

To determine how EBV orchestrates the cHL TME for immunomodulation and reorganization, we utilized Multiplexed Ion Beam Imaging (MIBI) spatial proteomics (18) and Nanostring GeoMx spatial transcriptomics (19) modalities to interrogate and delineate tumor-immune interactions within the native TME. We define distinct immune suppressive mechanisms in EBV-positive and EBV-negative cHL associated with different T-cell dysfunction states. These findings establish EBV as a critical modulator of the cHL TME, which may inform immunotherapeutic strategies in EBV-positive cHL and other viral-associated malignancies.

## Results

### Spatial Proteomics to Interrogate Differential TME Responses Between EBV-positive and EBV-negative cHL

We developed and applied a 30-plex Multiplexed Ion Beam Imaging (MIBI) antibody panel (**Supp Table 1**), including phenotyping and functional markers, to archival excisional formalin-fixed paraffin-embedded (FFPE) tumor tissues from cHL patients (6 EBV-positive, 14 EBV-negative; **Supp Table 2**), for downstream cell type annotation and cell neighborhood (CN) identification to dissect EBV-linked tumor-immune responses in the cHL TME (**Fig. 1A**). All antibodies were titrated and optimized in reactive lymph nodes and cHL cases (**Fig. 1B** and **Supp Fig. 1**) to ensure binding specificity and optimal signal-noise as previously described (20–22). We confirmed the panel specificity for cell phenotyping and functional assessment, including markers for M2-like macrophages (CD163), B cells (Pax5^hi^), HRS cells (Pax5^lo^), dendritic cells (CD11c), PD-L1 (**Fig. 1B**, top), and the localization of T-cell specific markers (CD4, CD8) and their memory (CD45RO) and dysfunctional (Tox) subpopulations (**Fig. 1B**, bottom). Examples of staining for all 30 markers are included in **Supp Fig. 1**.

**Figure 1.**
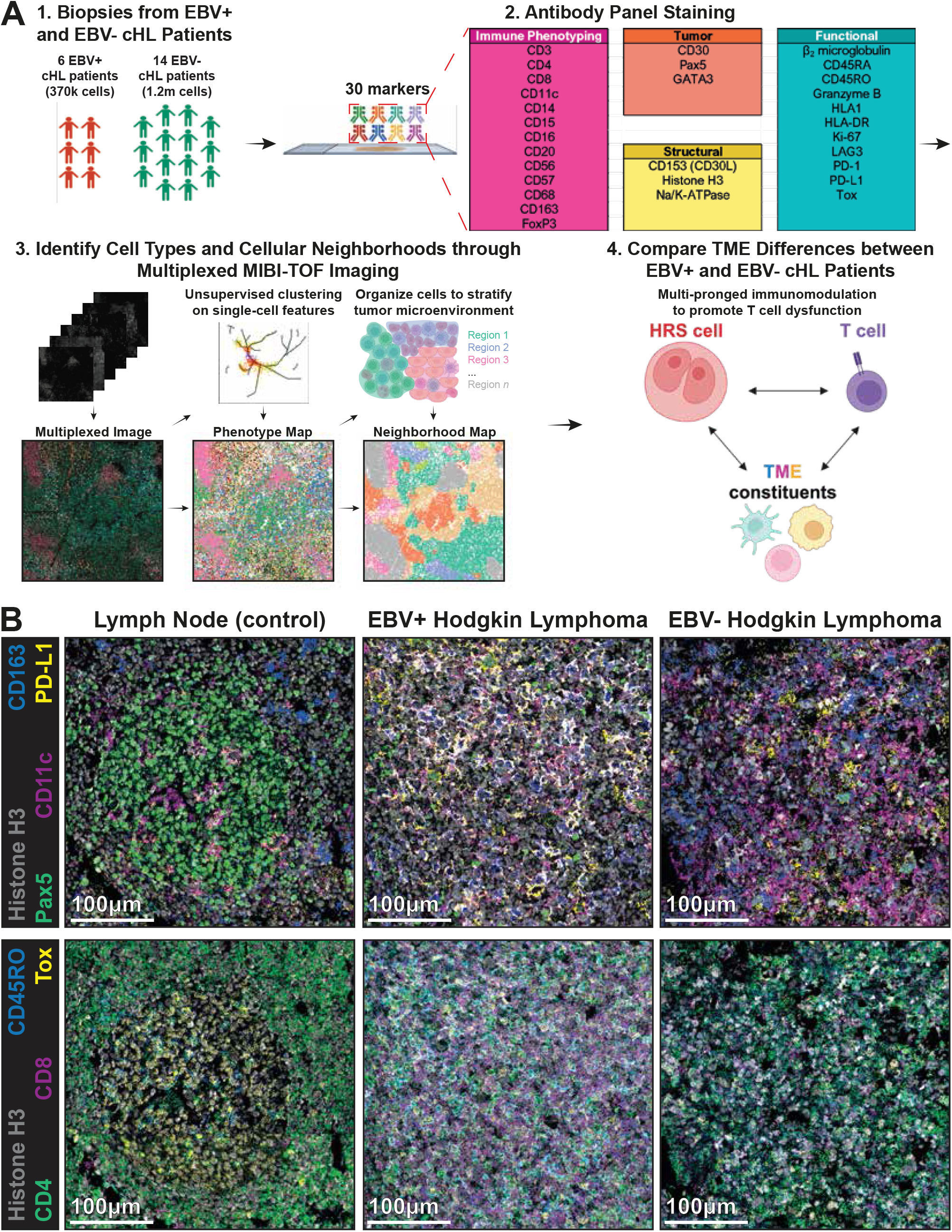
An overview of the spatial-omics framework to interrogate differences between EBV-positive and EBV-negative cHL TME. **(A)** Study overview to spatially dissect the EBV-positive and EBV-negative cHL TME. (1) Biopsies from an initial cohort of EBV-positive (n=6) and EBV-negative (n=14) cHL patients were collected, (2) stained with cocktail of 30 lanthanide-conjugated antibodies, and acquired by the MIBI spatial proteomics platform to (3) generate multiplexed images that allowed for the identification of cell phenotypes and neighborhoods, where upon (4) computational analysis illustrated key differences between EBV-positive and EBV-negative cHL TMEs. **(B)** Representative MIBI images of lymph node (left), EBV-positive (middle), and EBV-negative cHL (right) sections, with markers for nuclei (Histone H3), M2-like macrophages (CD163), HRS cells (Pax5^lo^), dendritic cells (CD11c), programmed death-ligand 1 (PD-L1), memory T cells (CD45RO), CD4 T cells (CD4), CD8 T cells (CD8), and T-cell dysfunction-associated transcription factor (Tox). Scale bar: 100 µm.

### Immune Components and Higher T-cell dysfunction Signatures Distinguish EBV-positive From EBV-negative cHL TME

We next performed cell segmentation (**Supp Table 3**) and clustering across all 20 patients, as previously described (20–22), over 41 stitched field-of-views (FOVs) comprised of 794 individual tiles of 400×400 µm images, as selected by a team of expert hematopathologists using ad-jacent H&Es (see **Materials & Methods**). All cell annotations (**Supp Fig. 2**) were stringently inspected and confirmed on the original MIBI multiplexed images for accuracy with expert hematopathologists. Key phenotypic markers were quantified to further confirm their expected enrichment on the corresponding cell types (**Fig. 2A**, top left and **Supp Fig. 3A**), with their functional markers (not used for clustering and cell type identification) depicted separately below (**Fig. 2A**, bottom left and **Supp Fig. 3A**). All annotated cells were evaluated via inspection of the original images, including confirmation of background due to lateral signal spillover, an issue common in spatial proteomics data (23), to confirm that “spillover” effects did not impact cell quantitation or other downstream analyses. For example, high CD3 (and some CD4) signals in dendritic cells (DCs, identified using CD11c) were confirmed to be background due to the close proximity of CD11c+ cells to CD3+ cells (**Supp Fig. 3B**, left), and PD-L1 signals on neutrophils (identified using CD15) was confirmed to be background due to their close proximity to PD-L1 expressing cells such as DCs (23) (**Supp Fig. 3B**, right).

**Figure 2.**
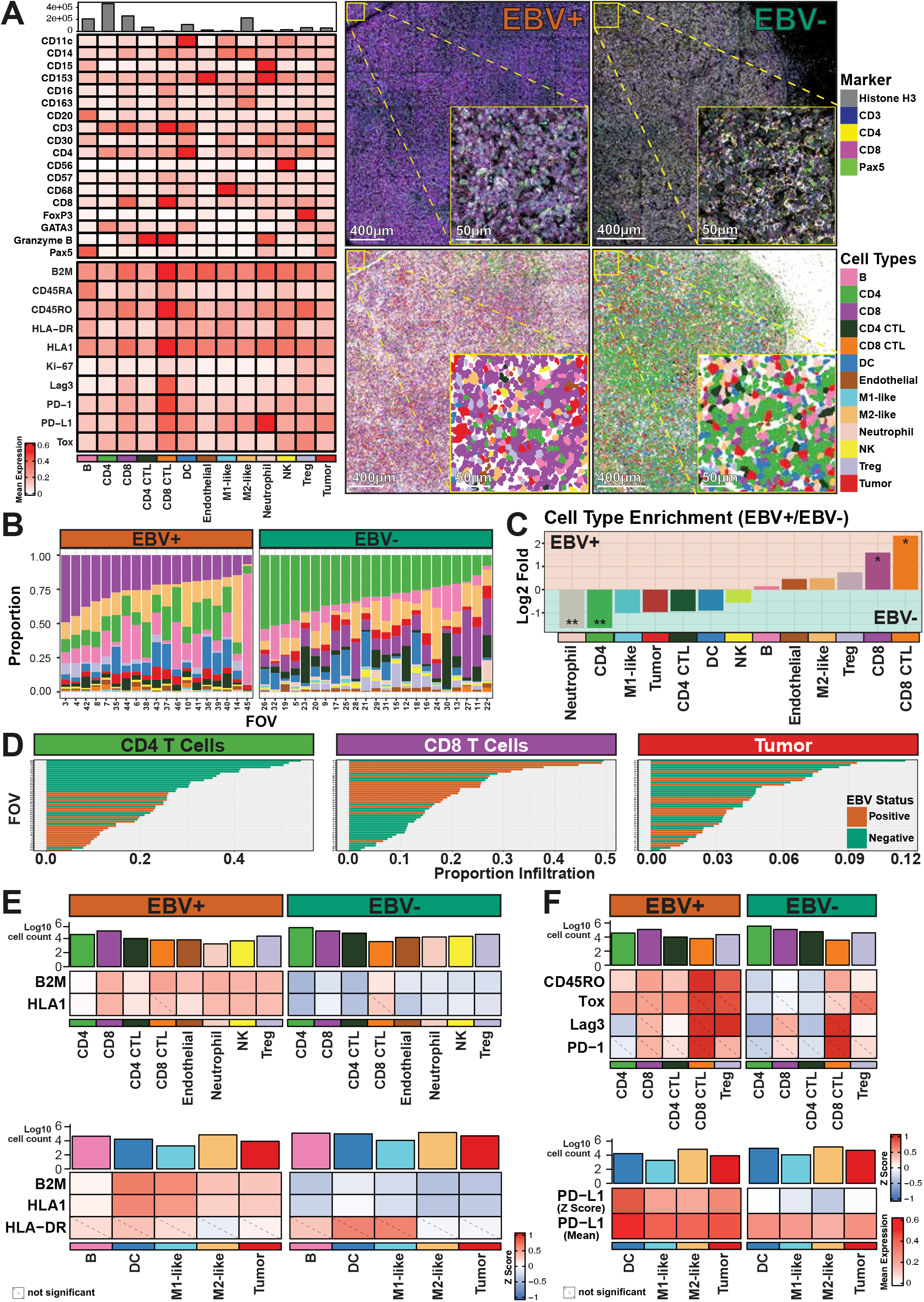
Distinct immune components and T-cell dysfunction features between EBV-positive and EBV-negative cHL TME. **(A)** Relative mean expression levels of phenotypic (left, top) and functional markers (left, bottom) for the annotated cell phenotypes in this MIBI dataset. A z-score visualization of this heatmap is in **Supp Fig. 3A**. Representative MIBI images of EBV-positive (middle, top) and EBV-negative cHL FOVs (right, top), with markers for nuclei (Histone H3), T cells (CD3), CD4 T cells (CD4), CD8 T cells (CD8), B cells (Pax5^hi^), and HRS cells (Pax5^lo^), along with the corresponding segmented and annotated EBV-positive (middle, bottom) and EBV-negative cell phenotype maps (right, bottom). Phenotype maps for the other FOVs are in **Supp Fig. 2**. Scale bar: 400 µm (main), 50 µm (inset). **(B)** Relative proportions of annotated cell types across EBV-positive and EBV-negative cHL FOVs. **(C)** Log2 fold enrichment plot of annotated cell types between EBV-positive and EBV-negative cHL FOVs. Significance stars (* *p ≤* 0.05, ** *p ≤* 0.01) are only shown for statistically significant comparisons (*p ≤* 0.05). Two-sided Wilcoxon rank sum tests were conducted for all cell type proportions, with alternative hypotheses that the proportions of the given cell type present in the two strata were not equal. Test results were adjusted for multiple comparisons using the Benjamini-Hochberg method with a targeted false discovery rate (FDR) at 0.05. Unadjusted p-value and Benjamini-Hochberg corrected test results are in **Supp Table 4. (D)** Relative proportion infiltration of CD4 T, CD8 T, and HRS cells across all FOVs. **(E)** Top: Relative expression of the MHC Class I components (B2M and HLA1) on annotated cell types, apart from antigen-presenting cells and HRS cells, in the cHL TME. Bottom: Relative expression of MHC Class I and MHC Class II (HLA-DR) on antigen-presenting cells and HRS cells in the cHL TME. Representative MIBI images of MHC Class I differences are in **Supp Fig. 3E. (F)** Top: Relative expression of memory (CD45RO) and T-cell dysfunction (Tox, Lag3, PD-1) markers on T-cell populations, with the relative cell counts inscribed above. Bottom: Relative expression of programmed death-ligand 1(PD-L1) on antigen-presenting cells (DCs and macrophages (M1-like, M2-like)) and HRS cells, with the relative cell counts inscribed above; both z-score and mean expression representations are here. Statistical test results for **(E) & (F)** are in **Supp Figs. 3B & 3E**, and unadjusted p-value and Benjamini-Hochberg corrected test results are in **Supp Table 5**. Mean expression visualization for **(E) & (F)** heatmaps are in **Supp Figs. 3C & 3F**.

**Figure 3.**
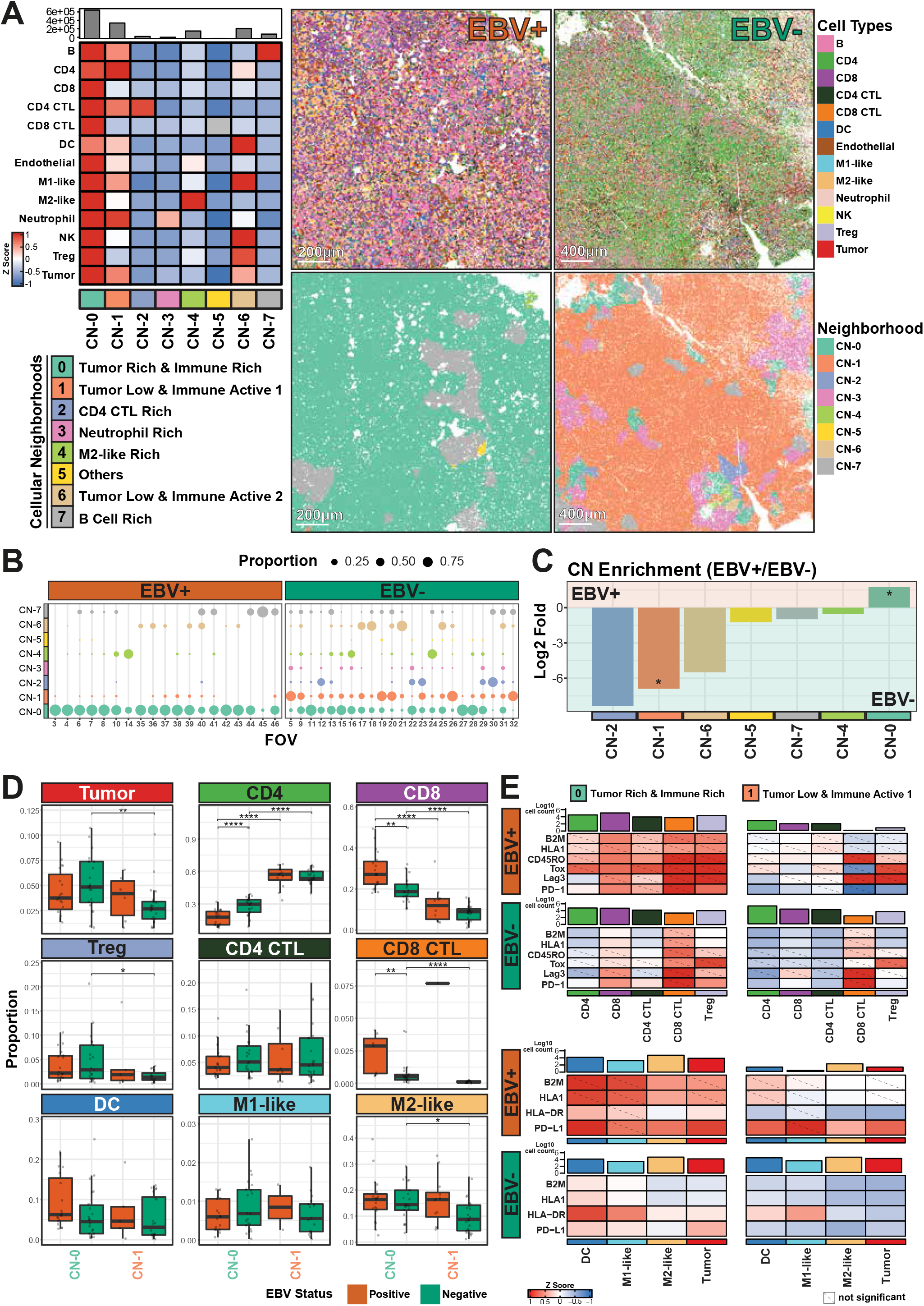
Cell neighborhood and EBV-linked T-cell dysfunction in the cHL TME. **(A)** Relative proportions of cell types (left) across the 8 identified cell neighborhoods (CN) in the EBV-positive and EBV-negative cHL MIBI dataset. Representative annotated cell phenotype maps of EBV-positive (middle, top) and EBV-negative cHL FOVs (right, top), along with the visualization of their corresponding EBV-positive (middle, bottom) and EBV-negative CNs (right, bottom). CN maps for the other FOVs are in **Supp Fig. 4**. Scale bar: 200 µm (Top, EBV+), 400 µm (Bottom, EBV-). **(B)** Relative proportion of cells assigned to each CN across EBV-positive and EBV-negative cHL FOVs. **(C)** Log2 fold enrichment plot of cell proportions assigned to each CN between EBV-positive and EBV-negative cHL FOVs. Note that CN3 is not shown due to its absence from the EBV+ cHL TME. Significance stars (* *p ≤* 0.05) are only shown for statistically significant comparisons (*p ≤* 0.05). Two-sided Wilcoxon rank sum tests were conducted for all cell type proportions within each CN, with alternative hypotheses that the proportions of the given cell type in each CN present in the two strata were not equal. Test results were adjusted for multiple comparisons using the Benjamini-Hochberg method with a targeted false discovery rate (FDR) at 0.05. Unadjusted p-value and Benjamini-Hochberg corrected test results are in **Supp Table 6. (D)** Relative proportion of the major immunomodulatory cell types identified in **Fig. 2F** within CN-0 and CN-1, and stratified by EBV status. Two-sided Wilcoxon rank sum tests were conducted for all combination of cell type, cell neighborhood, and EBV status, with alternative hypotheses that the proportions of the given cell type present in the two strata were not equal. Test results were adjusted for multiple comparisons using the Benjamini-Hochberg method with a targeted false discovery rate (FDR) at 0.05. Unadjusted p-value and Benjamini-Hochberg corrected test results are in **Supp Table 8**. Significance stars (* *p ≤* 0.05, ** *p ≤* 0.01, *** *p ≤* 0.001, **** *p ≤* 0.0001) are only shown for statistically significant comparisons after multiple comparisons correction. Data for the other cell types in the cHL TME are in **Supp Fig. 5A**, and unadjusted p-value and Benjamini-Hochberg corrected test results are in **Supp Table 7. (E)** Relative z-score expression level of markers for MHC Class I (B2M and HLA-1), MHC Class II (HLA-DR), memory T cells (CD45RO), and T-cell dysfunction features (Tox, Lag3, PD-1, PD-L1) on the relevant cell types with the relative cell counts inscribed above, stratified by CN (columns) and EBV status (rows). Statistical test results for **(E)** are in **Supp Fig. 5B**, and the unadjusted p-value and Benjamini-Hochberg corrected test results are in **Supp Table 8**. Mean expression visualization of this heatmap are in **Supp Fig. 5C**.

We observed relative differences in the TME composition between EBV-positive and EBV-negative cHL TMEs (**Fig. 2A**, right), and further quantified the differences in cell type proportions (**Fig. 2B**) and enrichment (**Fig. 2C** and **Supp Table 4**). Of note, there were significantly higher proportions of CD8 T cells in the EBV-positive TME, in stark contrast with the significant increase of CD4 T-cell and neutrophil proportions in the EBV-negative cHL TME (**Figs. 2A-C**). This result is also consistent when evaluated on individual FOV levels (**Fig. 2D**). Our data also revealed the prevalence of M2-like macrophages and Tregs in the EBV-positive TME (**Fig. 2C**), and observed a generally higher number of HRS cells in EBV-negative cHLs (**Fig. 2C**).

The presence of elevated CD8 T cells, coupled with their ineffective tumoricidal function, implicates EBV in uniquely conditioning the cHL TME to actively suppress T-cell effector responses. We tested this hypothesis by quantifying the average MHC Class I (**Fig. 2E**, B2M & HLA1 and **Supp Table 5**) and Class II (**Fig. 2E**, HLA-DR and **Supp Table 5**) expression on cells within the cHL TME across the imaged FOVs. Our results show significantly higher MHC Class I broadly across all cell types, including HRS cells, in the EBV-positive cHL TME, whereas differences in MHC Class II expression were not statistically significant (**Fig. 2E, Supp Fig. 3C**, and **Supp Table 5**). As all non-neoplastic cells express MHC Class I (and 50% of EBV-positive and 10% of EBV-negative HRS cells (15)), we re-visualized these z-score expression heatmaps with a mean expression heatmap (**Supp Fig. 3D**) to confirm the relative expression of MHC Class I across TME immune constituents, thus verifying that these z-score heatmaps emphasize only the differences in MHC Class I expression by EBV status. Visual reinspection of raw MIBI images also confirmed a tissue-level increase in MHC Class I expression in the EBV-positive cHL TME beyond HRS cells (**Supp Fig. 3E**). These results expand upon previous observations on differential Class I and II expression in EBV-positive and EBV-negative HRS cells (12, 13, 15, 16, 24), and highlight global antigen presentation differences in the TME across various cell types. The cHL TME is known to be significantly enriched in PD-L1 (24, 25), thus implicating PD1+ T cells as a key target of immunosuppression. We therefore further dissected memory (CD45RO (26)) and immune checkpoint regulators (Tox (27–30), Lag3 (31), PD-1 (32)) on T cells. Although dysfunctional CD8 and CD4 T cells have contrasting molecular profiles (33), both Lag3 and PD-1 are key features of CD8 and CD4 T-cell dysfunction (33, 34); Tox, which has an established role in CD8 T-cell dysfunction and a less clearly defined function in CD4 T cells, was also utilized an additional proxy for potential CD4 T-cell dysfunction. We found a significant increase in T-cell memory (CD45RO) and dysfunction (Tox, Lag3) markers in an EBV-linked manner on CD4 T cells, but not PD-1 (**Fig. 2F**, top, **Supp Fig. 3F**, and **Supp Table 5**). We additionally observed a significant increase in PD-L1, the ligand for PD-1, on both HRS cells and antigen-presenting cells (DCs, M1-like, and M2-like macrophages) in the EBV-positive TME compared to the EBV-negative TME (**Fig. 2F**, bottom, **Supp Fig. 3F**, and **Supp Table 5**). This was re-visualized with a mean expression heatmap (**Fig. 2F**, bottom) to confirm that HRS cells and antigen-presenting cells (DCs and macrophages (M1-like, M2-like)) have substantial PD-L1 expression (24, 25) regardless of EBV status; z-score expression heatmaps for T-cell activation and dysfunction markers were similarly re-visualized using mean expression heatmaps (**Supp Fig. 3G**) to confirm that T cells express dysfunction markers (35, 36) regardless of EBV status. Further stratification of T cells into memory and naive populations, based on the relative expression of CD45RO and CD45RA, revealed a consistent enrichment of dysfunction (Tox, Lag3, PD-1) and proliferation (Ki67) markers in memory T cells (**Supp Fig. 3H**), implicating previously activated T cells as a major source of T-cell dysfunction within the cHL TME.

These results expand upon previous observations regarding PD-1 and Lag3 expression on CD4 T cells and PD-L1 on macrophages within the immunoprivileged cHL TME (36) to include increased M2-like macrophage and Treg populations (**Fig. 2C**) in the EBV-positive cHL TME, and potentially suggest engagement of the PD-1/PD-L1 axis between T cells, HRS cells, and surrounding immune constituents to coordinate EBV-linked immune dysregulation of the cHL TME. Our observations of checkpoint modulation of T cells by HRS cells is consistent with a model in which EBV further exploits these pathways to dampen productive immune responses. These results suggests an EBV-dependent mechanism for suppressing T-cell effector functions through reorganization and re-education of the cHL TME.

### Tumor Cellular Neighborhood-specific T-cell Dysfunction and Antigen presentation in the cHL TME

We next computationally defined the cellular neighborhoods (CNs) responsible for spatial reorganization within the cHL TME by adopting the spatial latent Dirichlet allocation (LDA) approach (37) to systematically identify 8 CNs across both EBV-positive and EBV-negative cHLs (**Fig. 3A** and **Supp Fig. 4**). We observed that HRS cells are primarily located in CN-0, CN-1, and CN-6 (**Fig. 3A**), and these CNs comprise distinct T-cell populations. While CN-0 (Tumor Rich & Immune Rich) is diverse in T-cell and immune cell composition, CN-1 (Tumor Low and Immune Active 1) has a lower proportion of CD8 T cells, and CN-6 (Tumor Low and Immune Active 2) is generally deficient in T cells with the exception of Tregs. In addition, M2-like macrophages are also more enriched in CN-0 compared to CN-1 and CN-2 (**Fig. 3A**). Stratification by EBV status revealed that CN-0 was generally present across all cHL samples, but particularly dominant in the EBV-positive cHL patients, while a mixture of the other CNs, including CN-1 and CN-6, were prevalent in EBV-negative cHL TMEs (**Figs. 3B & C**). As HRS cells were most abundant in CN-0 followed by CN-1 (**Fig. 3A**), CN-0 is significantly enriched in EBV-positive TMEs, and CN-1 is significantly enriched in EBV-negative TMEs (**Fig. 3C** and **Supp Table 6**), we further analyzed the phenotypic and functional constituents in these two CNs to elucidate the key factors that selectively modulate tumor-immune interactions in the EBV-positive and EBV-negative cHL TME. We omitted CN-6 from this analysis due to its selective prevalence in a subset of EBV-positive and EBV-negative patients, whereas CN-1 and CN-0 were more generally present across both EBV-positive and EBV-negative TMEs (**Fig. 3B**), and were statistically significantly different in the CN proportion analysis (**Fig. 3C** and **Supp Table 6**).

**Figure 4.**
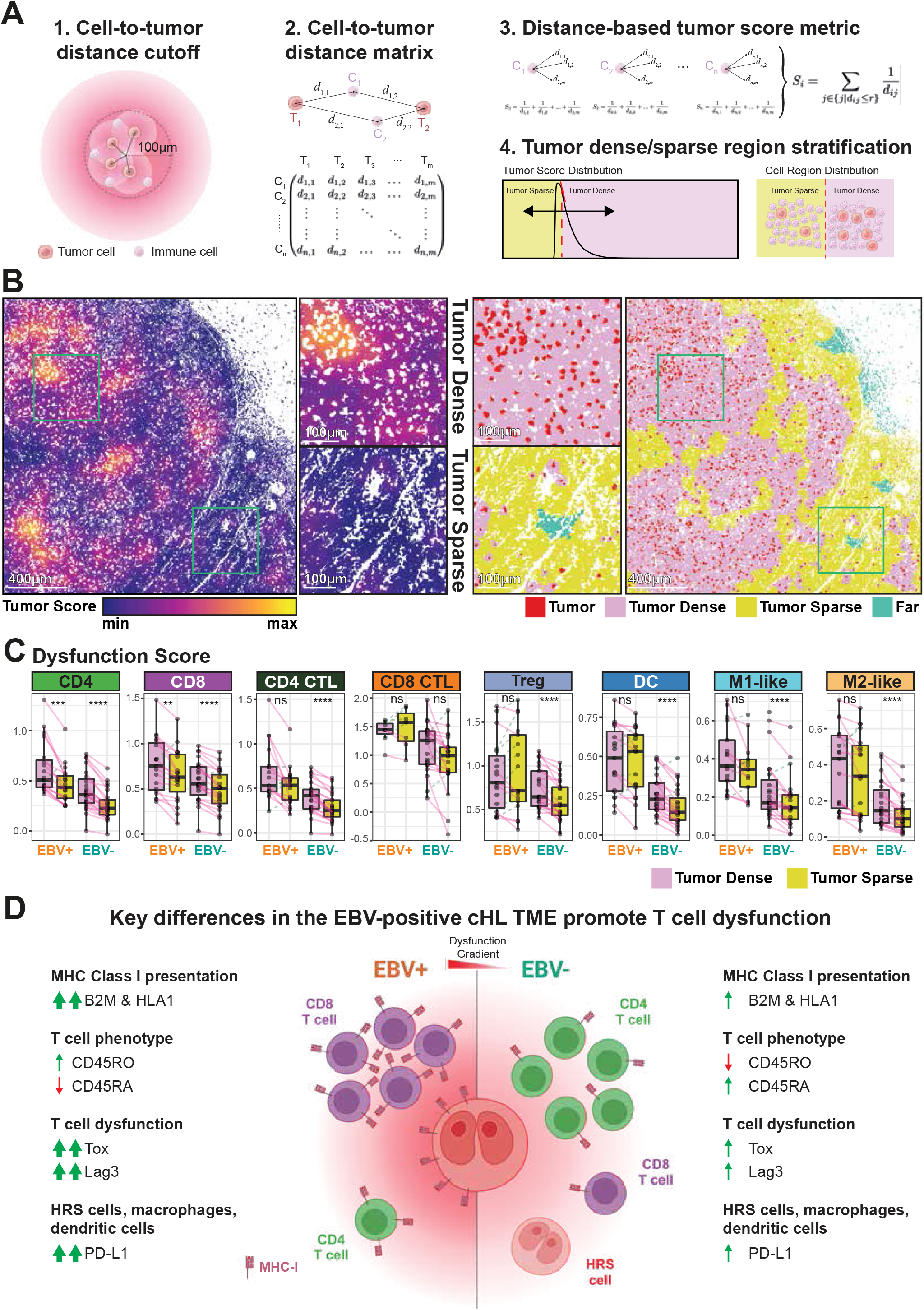
Immune Dysfunction in the cHL TME is a Function of Spatial Proximity to HRS cells. **(A)** Schematic of our approach to stratify cells into tumor-dense and tumor-sparse populations. (1) Cell-to-tumor distances for non-tumor cells within 100 µm from each tumor cell were (2) organized into a cell-to-tumor distance metric and used to (3) calculate the “tumor score” metric for each non-tumor cells; this “tumor score” metric was then used to (4) stratify cells into into tumor sparse and tumor dense regions based on the global tumor score distribution. **(B)** Representative tumor score map (left), along with the visualization of tumor dense and tumor sparse regions (right), in a cHL FOV. Tumor cells have a white mask in the tumor score map because they do not have a tumor score. Tumor score map and dense/sparse regions for all FOVs are in **Supp Fig. 6** and **7** respectively. Scale bar: 400 µm (main), 100 µm (inset). **(C)** Dysfunction scores of the major immunomodulatory cell types identified in **Fig. 2F** within tumor dense and tumor spare regions are stratified by EBV status, with each dot representing one FOV. For each tumor dense and tumor sparse pair, a one-sided T test was performed, with the alternative hypothesis that the difference between the mean dysfunction score of the tumor dense region in an FOV and that of the tumor sparse region is greater than 0 (* *p ≤* 0.05, ** *p ≤* 0.01, *** *p ≤* 0.001, **** *p ≤* 0.0001). The test results were adjusted for multiple comparisons using the Benjamini-Hochberg method with a targeted FDR at 0.05. The expression of individual markers used to generate the dysfunction score metric are in **Supp Fig. 8**, and the unadjusted p-value and Benjamini-Hochberg corrected test results are in **Supp Table 9. (D)** Cartoon depicting the key differences between EBV-positive and EBV-negative cHL TME.

We first focused our investigations on cellular compositional differences amongst HRS cells and immune components within these two HRS cell-enriched CNs. Stratification of CN-0 by EBV status revealed significant enrichment of CD8 and CD8 CTL in the EBV-positive TME and CD4 enrich-ment in the EBV-negative TME (**Fig. 3D** and **Supp Table 7**), in line with our previous data (**Figs. 2C & D**). We further observed, in CN-0, a consistent number of HRS cells, Tregs, DCs, macrophages (M1-like and M2-like), and endothelial cells in EBV-positive and EBV-negative TMEs, and an in-crease in B cell and decrease in neutrophil, and NK cell pop-ulations in the EBV-positive TME (**Fig. 3D, Supp Fig. 5A**, and **Supp Table 7**). We also observed a relatively consis-tent number of HRS cells and immune cell populations in CN-1, apart from a modest increase of M2-like macrophages in the EBV-positive TMEs (**Fig. 3D** and **Supp Fig. 5A**). The minor differences in the number of HRS cells between EBV-positive and EBV-negative TMEs within CN-0 and CN-1 (**Fig. 3D** and **Supp Table 7**) suggests that the quantity of HRS cells alone is not a key differentiator of EBV-linked dif-ferences in cHL TME immune dysfunction.

**Figure 5.**
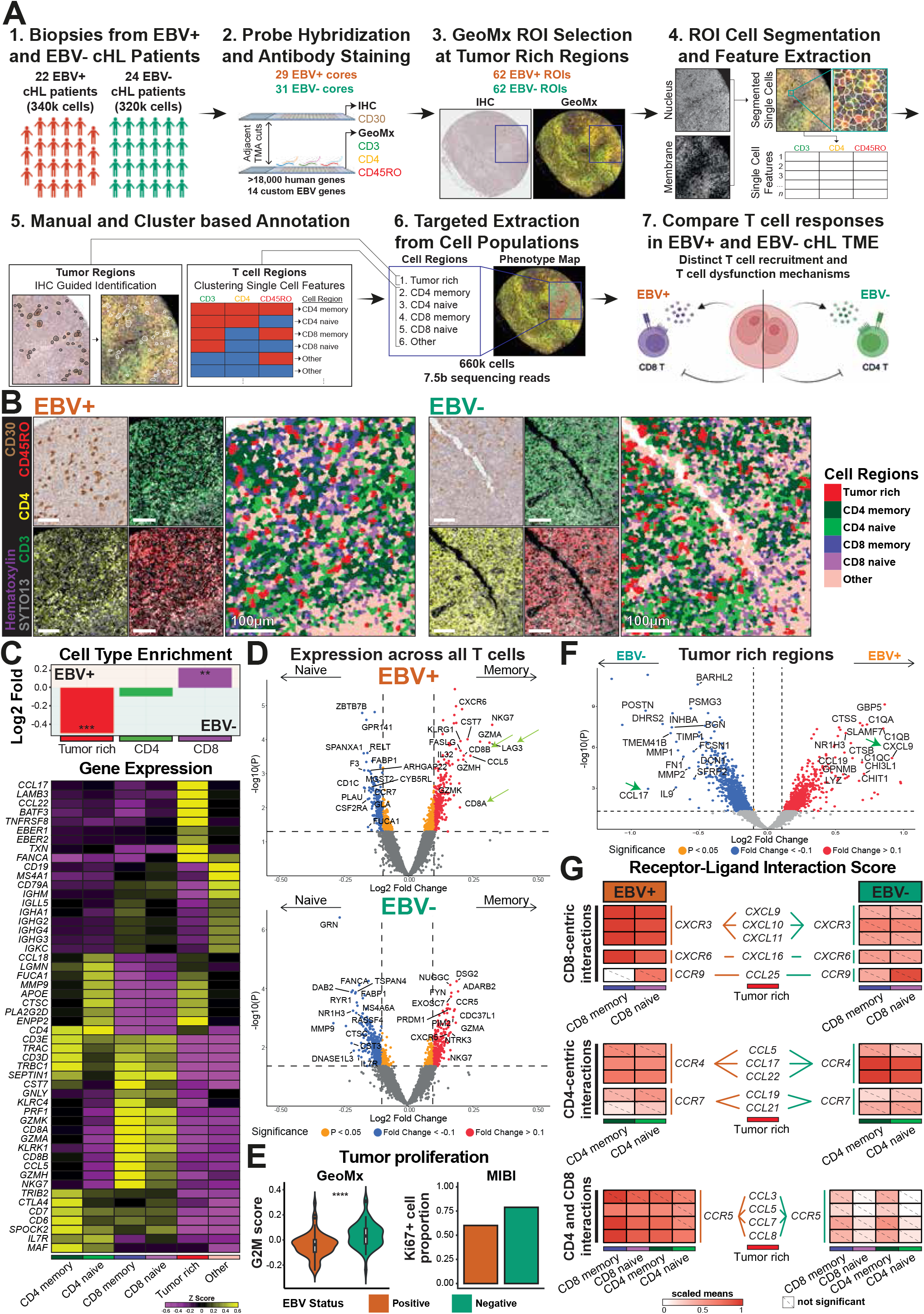
A spatial multi-omics framework reveals EBV-linked modulation of tumor proliferation and cytokine signatures within the EBV-positive cHL TME. **(A)** Our customized multi-omics workflow to evaluate EBV-positive and EBV-negative cHL TME. (1) Biopsies from two cohorts of EBV-positive (n=22) and EBV-negative (n=24) cHL patients were assembled into TMAs, where (2) one TMA section was stained with anti-CD30 to visualize HRS cells through IHC (top), and the adjacent TMA section hybridized with an >18,000 RNA probe panel targeting human and EBV genes and stained with three fluorescent antibodies for GeoMx assay (bottom). This allowed the (3) selection of 124 tumor-enriched ROIs (n=62 for both EBV-positive and EBV-negative cores), (4) computational identification of single cells and marker expression within each ROI, and (5) annotation of 6 different tumor, T, and immune cell populations, leading to (6) targeted extraction of wholte transcriptomes within each cell region that enable the (7) dissection of key differences underlying distinct T-cell populations and dysregulation between EBV-positive and EBV-negative cHL TMEs. **(B)** Representative IHC and immunofluorescent images of EBV-positive (left) and EBV-negative cHL (right) sections, with markers for HRS cells (CD30), CD8 T cells (CD3+ & CD4-), CD4 T cells (CD3+ & CD4+), memory T cells (CD45RO), and nuclei (Hematoxylin and SYTO13) shown in the smaller panels, and the corresponding cell annotations shown in the larger panels. Scale bar: 100 µm. **(C)** Top: Log2 fold enrichment plot of annotated cell regions between EBV-positive and EBV-negative cHL ROIs. Significance stars (* *p ≤* 0.05, ** *p ≤* 0.01, *** *p ≤* 0.001) are only shown for statistically significant comparisons (*p ≤* 0.05). Two-sided Wilcoxon rank sum tests were conducted for all cell type proportions, with alternative hypotheses that the proportions of the given cell type present in the two strata were not equal. Test results were adjusted for multiple comparisons using the Benjamini-Hochberg method with a targeted false discovery rate (FDR) at 0.05. Unadjusted p-value and Benjamini-Hochberg corrected test results are in **Supp Table 12**. Bottom: Expression heatmap of the key genes associated with each annotated cell region. Expression heatmap of other T-cell cytotoxic genes and EBV genes are respectively in **Supp Fig. 9A** and **9B. (D)** Volcano plots showing differences in gene expression between memory (CD45RO+) and naive T cells (CD45RO-) stratified by EBV status, with a few of the most differentially expressed genes indicated. Notice the green arrows that point to *CD8A, CD8B*, and *LAG3* transcripts. The volcano plot without EBV stratification is in **Supp Fig. 9C. (E)** Orthogonal validation of G2M cell proliferation transcriptomic signatures between EBV-positive and EBV-negative tumor regions from GeoMx cohort (*** *p ≤* 0.001) (left), and Ki67+ HRS cell proportion from MIBI cohort (right). A two-sided Wilcoxon rank sum test was conducted for the G2M signature, with the alternative hypothesis that the G2M signature in the two strata was not equal. Test results were adjusted for multiple comparisons using the Benjamini-Hochberg method with a targeted false discovery rate (FDR) at 0.05. No statistical tests were performed for the Ki67+ cell proportion as it shows only two discrete values. **(F)** Volcano plot of EBV-positive vs. EBV-negative tumor rich regions, with some of the highly differentially expressed host genes shown. *CXCL9* and *CCL17* are indicated by the green arrows. **(G)** Receptor-ligand analysis of chemokines that promote T-cell recruitment. Significance stars (* *p ≤* 0.05, ** *p ≤* 0.01, *** *p ≤* 0.001, **** *p ≤* 0.0001) are only shown for statistically significant comparisons (*p ≤* 0.05). P values were generated from one-sided permutation tests, with the alternative hypothesis that the distribution of a given receptor-ligand interaction is greater than the null distribution (see Materials & Methods).

Given the enrichment of PD-L1 in the cHL TME (24, 25), and our strong evidence for EBV-linked T-cell dysfunction (**Fig. 2F**), we next focused on a functional dissection of cellular states by comparing antigen presentation pathways, T-cell activation, and T-cell dysfunction states between CN-0 and CN1 (**Fig. 3E, Supp Figs. 5B & 5C**, and **Supp Table 8**). Our results reflect a global increase in MHC Class I expression (HLA1 and B2M), T-cell activation (CD45RO), T-cell dysfunction (Tox, Lag3, PD-1) regardless of CD8 or CD4 lineages, and higher expression of PD-L1 on HRS cells and antigen-presenting cells (DCs and macrophages (M1-like, M2-like)), predominant observations particularly for CN-0 and EBV-positive TMEs (**Fig. 3E, Supp Fig. 5B**, and **Supp Table 8**). We similarly confirmed that these features were all expressed across both CN-0 and CN-1 by re-visualizing these z-score expression heatmaps with mean expression heatmaps (**Supp Fig. 5C**). These observations are consistent with a model in which HRS cells exert a direct influence within their immediate neighborhoods to condition the cHL TME and promote T-cell dysreguation regardless of EBV-status, but EBV-positive HRS cells appear to additionally promote MHC-I antigen presentation and further dysregulate T cells.

### Identification of Immune Dysfunction as a function of Spatial Organization around HRS cells in the cHL TME

Our results suggest that HRS cells may modulate immune dysfunction and suppression within their immediate microenvironment (**Figs. 3D & E**). We thus hypothesized that dysfunction of immune cells within the cHL TME is dependent on distance from their proximal HRS cells. We tested this by first developing a “Tumor Score” metric, which takes into account both the distance and density of nearby tumor cells from each immune cell of interest (within 100 µm) (**Fig. 4A**). We then computed the Tumor Score across all the cells in this study and stratified cells into “Tumor Dense” or “Tumor Sparse” categories based on their location on an automatic cut-off point on the global tumor score distribution (**Figs. 4A & B** and **Supp Figs. 6-7**, see **Materials & Methods** for more details).

**Figure 6.**
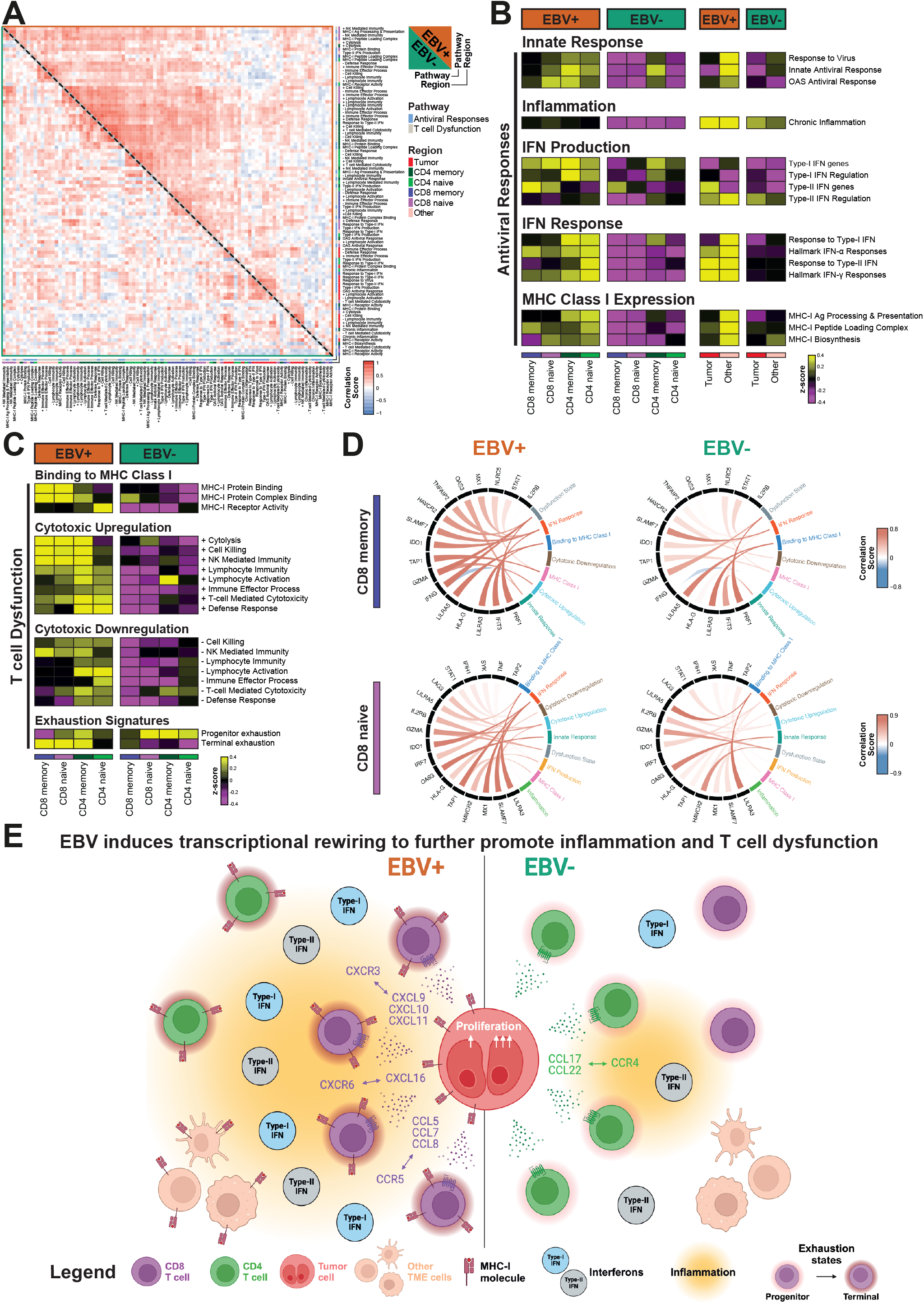
A systems-level functional dissection of the EBV-linked TME. **(A)** Spearman correlation heatmap of gene set enrichment scores across relevant cell regions between EBV-positive and EBV-negative cHL TME. Original names for the gene sets are in **Supp Table 13. (B)** Heatmap of gene set enrichment scores for pathways associated with antiviral responses, stratified by EBV status and cell region. Original names for the gene sets are in **Supp Table 13**, and expression of interferons associated with “Type-I IFN genes” and “Type-II IFN genes” pathways are in **Supp Fig. 10A. (C)** Heatmap of gene set enrichment scores for pathways associated with T-cell dysfunction, stratified by EBV status and cell region. Original names for the gene sets are in **Supp Table 13**, and the expression of genes associated with “Terminal exhaustion” and “Progenitor exhaustion” are in **Supp Fig. 10B. (D)** Circos plot displaying the Spearman correlation between the targeted pathway’s GSVA activity and corresponding gene expression. The black nodes represent genes, the colored nodes represent pathways, and links connecting pathways and genes are color-scaled by correlation coefficient. Circos plots for the other cell types are in **Supp Fig. 11. (E)** Our proposed model of how EBV orchestrates reorganization of the cHL TME.

We next assigned every immune cell a “Dysfunction Score”, placing a negative weight on the expression of markers linked to cell activation potential (CD45RA and Ki-67), and a positive weight on markers linked to T-cell activation and dysfunction (CD45RO, Tox, PD-1, and Lag3 for T cells, and PD-L1 for HRS cells and antigen presenting cells (DCs and macrophages (M1-like, M2-like) (see **Materials & Methods**) (34). We quantified the dysfunction scores of immune cell types in tumor dense and tumor sparse regions based on their Tumor Scores, to reveal a significantly higher dysfunctional state in tumor dense regions for both EBV-positive and EBV-negative cHL TMEs (**Fig. 4C, Supp Fig. 8**, and **Supp Table 9**). This increase in dysfunction score was notably observed across both CD8 and CD4 T cells. These results suggest that HRS cells play a pivotal role at modulating the cell states of immune constituents within the cHL TME. These proximity-linked dysfunction scores were also consistently elevated in EBV-positive regions compared to EBV-negative counterparts (**Fig. 4C** and **Supp Fig. 8**).

Our cumulative data supports a model in which EBV further enhances T-cell dysfunction and induces reorganization of the cHL TME distinctive from EBV-negative cHL (**Fig. 4D**). We highlight a tissue-level increase in MHC Class I antigen presentation (B2M and HLA1) and higher proportion of dysfunctional memory T cells of the CD8 lineage in the EBV-positive cHL TME, in contrast to the enriched CD4 T cells and higher density of HRS cells in the EBV-negative cHL TME. The proximity-linked immunomodulation of immune cells in the vicinity of HRS cells further promotes T-cell dysfunction, which is exacerbated through EBV-linked increase of markers including Tox, Lag3, PD-1, and PD-L1, potentially through direct chromatin regulation of these genes by EBV transcription factors (38).

### Dissection of Immune and HRS cells in the EBV-linked cHL TME Through Spatial Multi-omics on a Multi-institutional Cohort

As T-cell dysfunction is most pronounced when close to HRS cells (**Fig. 4C**), we next implemented a spatial multi-omics approach to further refine HRS cell and T-cell interactions leading to T-cell dysfunction in the cHL TME. Leveraging upon a multi-institutional cohort (22 EBV-positive, 24 EBV-negative; **Supp Table 2**), we performed GeoMx spatial genome-wide transcriptomic profiling (19) on 62 EBV-positive region-of-interest (ROI) and 62 EBV-negative ROIs (selected at areas dense in HRS cells) across 6 different cell populations (tumor rich, CD4 naive, CD4 memory, CD8 naive, CD8 memory, and Others) (**Fig. 5A**, see **Materials & Methods**). These 6 cell populations were identified using a combinatorial antibody staining (**Supp Table 10**), single cell segmentation (**Supp Table 3**), and clustering approach (**Supp Table 11**), with representative antibody staining patterns and cell type annotations shown in **Fig. 5B**. Given the design for the GeoMx in capturing transcripts in tissue regions and not at single-cell resolutions, as well as the potential presence of some T effector memory cells that re-express CD45RA (Temra) (39), we sought to verify if our defined regions were enriched for the denoted cell populations. We first confirmed that the enrichment of CD8 T cells in the EBV-positive cHL TME, in contrast to elevated CD4 and tumor cells in the EBV-negative cHL TME in this new cohort (**Fig. 5C**, top and **Supp Table 12**) was consistent with our MIBI results above (**Fig. 2C**, top). We next confirmed both the specificity of the assay and the accuracy of our cell type annotations via the top differentially expressed genes across the identified cell populations, including the enrichment of *TNFRSF8* (encoding CD30, a HRS marker) and the EBV-encoded small RNAs *EBER1* and *EBER2* in the Tumor Enriched region, CD4 and CD8 T-cell specific transcripts (such as *CD3D, CD3E, CD4, CD7, CD8A, CD8B*) in their respective regions, cytotoxic granzyme and perforin transcripts associated with T-cell effector functions (including *PRF1, GZMK, GZMA, GZMH*; more in **Supp Fig. 9A**) in memory T-cell regions, and immunoglobulin and B cell transcripts (including *CD19, MS4A1* (encoding CD20), *CD79A, IGH*s, *IGL*s, *IGK*s) found on B cells (under the “Other” region) (**Fig. 5C**, bottom). We also confirm the specific detection of EBV genes included in our customized EBV probe panel for the EBV-positive cHL tissues, including *EBER1, EBER2*, and *LMP1* as anticipated for EBV latency II in cHL, but also latent and lytic genes such as *EBNA2, BALF1, BCRF1, BNLF2A, BNLF2B, BZLF1*, and *RPMS1* (**Supp Fig. 9B**), consistent with bulk transcriptomic results from others (40). Our results demonstrate the robustness of the appropriate enrichment of genes within the 6 cell types we identified for spatial transcriptomics capture.

Differential gene expression analysis between memory and naive T cells, as defined by CD45RO expression, revealed an enrichment of *LAG3* transcripts in memory T cells (**Supp Fig. 9C**), in agreement with our MIBI findings (**Supp Fig. 3H**); Lag3 is an exhaustion marker for both CD8 and CD4 T cells (33). Further stratification of memory vs. naive comparisons by EBV status revealed the enrichment of *CD8A, CD8B*, and *LAG3* transcripts in memory T-cell populations in EBV-positive tissues (**Fig. 5D**, top), with no clear differences in T cell or dysfunction-linked transcripts between memory and naive T cells in EBV-negative tissues (**Fig. 5D**, bottom); this is consistent with the protein expression findings from our MIBI cohort, where T cells in EBV-positive tissues were predominantly memory CD8 T cells and exhibit greater features of immune dysfunction than those in EBV-negative tissues (**Figs. 2F & 3E**). We further note the elevated cytotoxic granzyme transcripts (including *GZMA, GZMH*, and *GZMK*) found primarily in the memory T-cell populations (**Fig. 5D**), in support of attempted effector functions within memory-differentiated T cells in the cHL TME. These transcriptomic-based results further revealed molecular details in support of our proposed model derived from spatial proteomics analysis (**Fig. 4D**).

We next focused on transcriptomic analysis of EBV-linked differences in tumor, CD8 T, and CD4 T-cell populations (**Fig. 5C**, top and **Fig. 2C**). We first orthogonally associated the EBV-linked decrease in tumor density to lower G2M cell cycle pathway activity in EBV-positive HRS tumors (**Fig. 5E**, left), consistent with the lower proportion of Ki67+ HRS cells in our MIBI results (**Fig. 5E**, right). Differential gene expression analysis between EBV-positive and EBV-negative tumor-rich regions further revealed contrasting enrichment of *CXCL9* (known to preferentially recruit CD8 T cells (41)) in EBV-positive regions and *CCL17* (shown to recruit CD4 T cells (42, 43)) in EBV-negative regions (**Fig. 5F**). We next performed a receptor-ligand interaction analysis (44, 45) between T cell and tumor regions within each ROI, for chemokine receptors that differentially modulate the homing of CD8 and CD4 T cells described within the contexts of cancer (41–43, 46–48). The EBV-positive cHL TME revealed pronounced receptor-ligand engagement for chemokines favoring CD8 T-cell attraction, including the *CXCR3*-*CXCL9*/*10*/*11* (41) and *CXCR6*-*CXCL16* (46) signaling axes (**Fig. 5G**, top panel), whereas those engaging CD4 T cells, such as the *CCR4*-*CCL17*/*22* axis (42, 43), were detected in EBV-positive and EBV-negative cHL TME, albeit at lower levels in the EBV-positive TME (Fig. 5G, middle panel). Our results also suggest an increase in engagement of the *CCR5*-*CCL5*/*7*/*8* receptor-ligand pairs (47, 48) specific to EBV-positive HRS and CD8 T-cell populations (**Fig. 5G**, bottom panel).

### Systems-level Inflammatory responses and Terminal Dysfunction states in the EBV-positive cHL TME

The elevated CD8 T-cell infiltration and MHC Class I presentation in an EBV-linked manner within the cHL TME are suggestive of increased antiviral and inflammatory immune responses against chronic viral persistence. We thus computed a global correlation for representative gene sets broadly categorized under “Antiviral Responses” and “T-cell dysfunction” separately for the EBV-positive and EBV-negative spatial transcriptomics data (**Supp Table 13**). We identified increased and orchestrated pathways enriched within the EBV-positive TME (**Fig. 6A**), suggesting a unique EBV-linked immunological landscape on the systems-level within cHL. We further delineated the enrichment scores of these associated gene sets stratified by cell population and EBV status to identify a tissue-level increase in immune responses linked to an antiviral state, including innate cellular immune responses, chronic inflammation, secretion and response to both Type-I and Type-II interferons (**Supp Fig. 10A**), and the expression of MHC Class I (**Fig. 6B**). The role of these pathways in promoting the expression of MHC Class I (49) implicates them in our observations of tissue-level increase in MHC Class I (**Fig. 2E**).

Our cumulative data and prior results (50) implicate T-cell dysfunction likely through unique EBV-linked mechanisms. We observed that although T cells in the EBV-positive cHL TME have higher expression of molecules that promote binding to MHC Class I and a paired increase in cytotoxic pathways, they also involved an increase in pathways that dampen cytotoxic responses (**Fig. 6C**). This contrasts the EBV-negative TME, in which antigen presentation and cytotoxicity were generally dampened.

Given our observation of HRS cell-dependent T-cell dysfunction in the cHL TME that was more pronounced in the presence of EBV (**Fig. 4C**), we sought to contextualize our results with the fact that T-cell exhaustion states exist across a progenitor-terminal continuum (34). We compared the expression of key genes that were identified to be differentially expressed between progenitor and terminally exhausted CD8 T-cells (51). While progenitor and terminal exhaustion remains under-investigated for CD4 T cells, we applied the same progenitor and terminal exhaustion transcriptional signatures on CD4 T cells as a proxy for CD4 T-cell exhaustion state. Interestingly, we found that T cells in the EBV-positive TME had a stronger terminally-differentiated exhaustion signature while those in the EBV-negative TME had a stronger progenitor-differentiated exhaustion signature regardless of CD8 or CD4 T-cell lineages (**Fig. 9C** and **Supp Fig. 10B**). These data suggest that EBV leads to extensive T-cell exhaustion in the cHL TME in a distinctive mechanism from their EBV-negative counterparts. Beyond T-cell exhaustion, we also noted an increase in *KLRG1* and *B3GAT1* (encoding CD57) expression and a decrease in *CD28* expression (**Supp Fig. 10C)**, suggestive of potential T-cell senescence and other T-cell dysfunction mechanisms (52) unique to EBV-positive cHL.

We finally sought to functionally link T-cell antiviral-associated responses with dysfunction in the EBV-stratified cHL TME (**Figs. 6B & 6C**) by visually linking the correlation between gene sets and their corresponding gene expressions for each cell type. We observed a global increase in T-cell and tumor-specific responses in CD8 T cells in an EBV-linked manner, as shown in a few of these representative genes that contribute to the “Antiviral Responses” and “T-cell dysfunction” pathways (**Fig. 6D**). These EBV-linked cell state differences were also consistently observed across other cell populations within the cHL TME (**Supp Fig. 11**), indicating that the presence of EBV can lead to tissue-level cell state transitions.

Our results support a model in which EBV induces HRS cells to condition their immediate microenvironment, including lower tumor proliferation and altered expression of chemokines and immunomodulatory ligands, for a CD8-heavy and highly dysfunctional T-cell landscape (**Fig. 6E**). The chronic viral persistence provokes an inflammatory response, including increased engagement with MHC Class I and interferon pathways, and an activated T-cell effector state that is further limited in effective tumoricidal programs through terminal exhaustion.

## Discussion

Establishment of an immunosuppressive TME is a central hallmark of cancer, one that is particularly prominent within the diverse immune milieu surrounding malignant HRS cells in cHL (1). Various mechanisms of immunosuppression in the cHL TME have been described, including the acquisition of somatic mutations and genomic rearrangements in HRS cells that lead to the frequent downregulation of MHC Class I and and less frequent dowregulation of MHC Class II pathways, and overexpression of PD-L1 and PD-L2 checkpoint ligands (14, 15, 24, 53, 54). Despite these advances, immunosuppression within the context of EBV-linked cHL and other tumors remains poorly understood, with EBV-positive HRS cells having a significantly lower mutational burden compared to their EBV-negative counterparts (54). We address this key knowledge gap by illustrating immune reorganization and immune suppression of the cHL TME in an EBV-linked fashion, with clear downstream implications. Our data is consistent with known features of the cHL TME, including the diversity of immune infiltrates such as lymphocytes, dendritic cells, macrophages, neutrophils, and natural killer cells (**Fig. 2A**), low density of HRS cells and general abundance of T cells (**Fig. 2A**) (55), high expression of PD-L1 on HRS cells and macrophages (**Figs. 2A & 2E**) (24, 54, 56–58), retention of MHC Class I expression in EBV-positive HRS cells (**Fig. 2E**) (12, 13), and evidence of potential PD-1/PD-L1 engagement (**Figs. 2F & 3F**) that attests to favorable patient responses to anti-PD-1 checkpoint immunotherapy (59).

One striking result from our study is that we found T cells in tumor-rich EBV-positive cHL tissues were predominantly of CD8 origin, memory-differentiated, and enriched in both effector and terminal exhaustion states. This represents an unusual T-cell dysfunction associated with EBV infection, and is supported by a recent single-cell RNA and T-cell receptor sequencing study that identified CD8 T-cell populations in the cHL TME as clonally expanded with effector and exhaustion transcriptional phenotypes (50). We therefore propose the following mechanism that leads to this T-cell dysfunction state. As the abundance of effector memory-differentiated CD8 T cells is coupled with a tissue-level increase in MHC Class I expression (HLA1, **Figs. 2E & 3E**) (suggestive of productive T-cell receptor engagement in EBV-positive TME), it is conceivable that these are elements of the host derived antiviral immune response, which includes antiviral interferon cytokines (potent inducers of MHC Class I (60)) (**Fig. 6B**), and increase in CD8 T-cell chemotaxis (**Fig. 5G**) into the TME. Consequently, the EBV-positive tumor cells counteracts this activated immune response by immunomodulation of the T cells into a terminally exhaustive state in a distance-dependent manner (**Figs. 2F, 3E, 4D**, & **6C**).

Due to the paucity of HRS cells, it is plausible that the cHL TME engages other immune constituents to maximize interactions with T cells and effectively subvert antitumor T-cell effector responses. It is noteworthy that other TME immune constituents, such as antigen-presenting cells (DCs and macrophages) that comprise a significant proportion of cells in the cHL TME (**Figs. 2A & 2B**), appear to participate in suppressing T-cell responses. This can be seen in part through the engagement of the PD-1/PD-L1 axis (**Figs. 2F & 3E**) due to PD-L1 upregulation in a distance-linked fashion from HRS cells (**Fig. 4C** and **Supp Fig. 8**), a potential mechanistic basis that warrants a deeper investigation into combinatorial efficacies of PD-1 based immunotherapies for the treatment of cHL (59). While checkpoint immunosuppression by HRS cells and macrophages through PD-1/PD-L1 has been described in cHL (24, 25), it has not been described for dendritic cells, thus identifying another potential cellular target involved in the effectiveness of checkpoint immunotherapies (61). Mechanisms of EBV-linked PD-L1 upregulation in antigen-presenting cells (DCs and macrophages) (**Fig. 4C** and **Supp Fig. 8**) will also be of significant interest given that they are not direct targets of EBV infection, but are potentially associated with the tissue-level increase in IFN-γ (**Fig. 6B** and **Supp Fig. 10A**) that is known to also promote PD-L1 expression (62). The higher expression of PD-L1 on EBV-positive compared to EBV-negative HRS cells (**Figs. 2F & 3E**) may be explained by both the increase in IFN-γ (62) (**Fig. 6B** and **Supp Fig. 10A**) and the expression of EBV latent oncoproteins LMP1 and EBNA2 (**Fig. 5D** and **Supp Fig. 9B**). LMP1 has been shown to promote PD-L1 expression in HRS cells by upregulating the activity of PD-L1 promoter elements (57) and its downstream NF-kB subunits (38); EBNA2, while more commonly associated with latency III, has been detected in HRS cells from RNAseq datasets (40) and is observed to be bound to enhancers directly linked to PD-L1 and PD-L2 genes (38). In addition to the potential increase in PD-1/PD-L1 engagement in the EBV-positive cHL TME (**Figs. 2-4**), T-cell populations in the EBV-positive TME appear to be more terminally exhausted (**Fig. 6C** and **Supp Fig. 10B**). Terminally exhausted T cells are known to have poorer tumor control and worse outcomes to PD-1 blockade compared to progenitor exhausted T cells, and terminal exhaustion can arise in lieu of chronic viral infection (51). However, T cells in the cHL TME have been reported to retain effector functions upon *ex vivo* restimulation regardless of severity of T-cell exhaustion (50), suggesting that T cells are not irreversibly dysfunctional in the EBV-positive cHL.

Limitations of this study include the lack of *in vitro* or *in vivo* models for functionally linking EBV infection in HRS cells with tissue-level TME reorganization. This is hindered by the lack of suitable murine models (63, 64), difficulties in establishing cHL lines that properly recapitulate their primary counterparts (65), and lack of organoid or organ-onchip models. Future advancements in these regards will be pivotal in addressing these limitations and instrumental in untangling EBV protein-specific functions in TME remodeling, currently beyond the scope of this study. Another limitation is the technical challenge to precisely categorize patients into those with HRS cells that express MHC Class I, and those with HRS cells that do not. This is in part due to the dispersed localization of HRS cells nested within immune cells that ubiquitously express MHC Class I (see B2M and HLA1 staining patterns in **Supp Fig. 1** and **Supp Fig. 3E**), the latter contributing to noise in adjacent segmented cells due to lateral signal spillover (23). As MHC Class I expression is retained in around 50% of EBV-positive and 10% of EBV-negative HRS cells (15), this limits a clearer mechanistic link between HRS cell MHC Class I status and T-cell dysfunction state that may be better evaluated with more sensitive and higher-resolution spatial-omics approaches. Nevertheless, our tissue-level analyses on CNs (**Fig. 3**) and tumor dense/sparse regions (**Fig. 4**) that are orthogonally validated on a transcriptomic level (**Figs. 5 & 6**) indicate that EBV-linked mechanisms, beyond the potential effects induced by MHC Class I status on HRS cells, are critical at promoting T-cell dysfunction. Furthermore, T-cell dysfunction is increasingly understood in the context of CD8 T cells but remains poorly known in CD4 T cells; our description of CD4 T-cell dysfunction relies on the assumption that these protein markers (Tox, Lag3, PD-1) and transcriptional signatures (**Supp Fig. 10B**, (51)) are consistent between dysfunctional CD8 and CD4 T cells, since both T-cell lineages have considerable overlaps in dysfunction signatures (including PD-1 and Lag3) despite having contrasting molecular profiles (33, 34). Our spatial multi-modal framework encompasses an initial discovery effort via spatial proteomics (**Figs. 1-4)**, followed by orthogonal validation and mechanistic dissection through spatial transcriptomics (**Figs. 5 & 6**), allowing a detailed investigation of tumor-immune interactions even when limited to archival clinical samples. This framework can be adapted for future studies of other diseases, highlighting both biological and technical breakthroughs.

In conclusion, we present here a detailed and iterative dissection of the tumor-immune interactions within the intact cHL TME. Our results contextualize prior findings with increased detail, providing new insights on the unique T-cell infiltration and cell states in a EBV-linked manner. Our results also highlight the need for technology-driven approaches to enable further biological insights, including the need to rethink the uniform clinical treatment of tumor virus-positive and - negative diseases.

## Materials & Methods

### Human tissue acquisition and patient consent

For MIBI analysis, formalin-fixed paraffin-embedded (FFPE) excisional biopsies from 20 patients with newly diagnosed cHL, and one reactive lymph node were retrieved from the archives of Brigham and Women’s Hospital (Boston, MA) with institutional review board approval (IRB# 2010P002736). All tumor regions were annotated by K.W., V.S. and S.J.R. For GeoMx spatial transcriptomics analysis, TMAs from two institutions were used. The Dana-Farber Cancer Institute TMA was constructed by S.S. and S.J.R. (IRB# 2016P002769 and 2014P001026), and includes 1 or 2 cores from each of 10 EBV-positive and 13 EBV-negative patients, as well as one tonsil control core, with each core measuring 1.5 mm in diameter. The University of Rochester Medical Center TMA was constructed by P.R. and W.R.B. (IRB# STUDY159) , and includes 1 core from each of 12 EBV-positive and 11 EBV-negative patients, as well as one tonsil control core, with each core measuring 2.0 mm in diameter. Tissues were sectioned onto gold slides (see below for more information) for MIBI analysis and SuperFrost glass slides (VWR, 48311-703) for GeoMx analysis, with each section measuring 5 µm in thickness. As part of the routine clinical pathology process, all cHL biopsies were confirmed to have HRS cells by immunohistochemical staining for CD30, and EBV status verified using *in situ* hybridization for *EBER*. Detailed EBV status for each patient is reported in **Supp Table 2**.

### MIBI Antibody conjugation

Antibody conjugation was performed according to a modified version of a previously published protocol (66). Maxpar X8 Multimetal Labeling Kit (Fluidigm, 201300) and Ionpath Conjugation Kits (Ionpath, 600XXX) were utilized. Briefly, 100 µg of BSA-free antibody was washed with the conjugation buffer, and then incubated with 4 µM TCEP (Thermo Fisher Scientific, 77720) for 30 minutes in a 37 °C water bath to reduce the thiol groups for conjugation. The reduced antibody was subsequently incubated with Lanthanide-loaded polymers for 90 minutes in a 37 °C water bath. The resulting conjugated antibody was purified by washing for five times with an Amicon Ultra filter with 50 kDa NMWL (Millipore Sigma, UFC505096). The conjugated antibody was quantified in IgG mode at A280 using a NanoDrop (Thermo Scientific, ND-2000). The final concentration was adjusted by adding at least 30% v/v Candor Antibody Stabilizer (Thermo Fisher Scientific, NC0414486) with additional 0.2% sodium azide, and the antibody was stored at 4°C.

### MIBI and GeoMx Antibody panel titration and validation

The antibody candidates used for cHL sample staining contain previously validated antibody clones (21, 67) and titers can be found in **Supp Table 1**, along with conjugated channels. In brief, antibody candidates were first validated for specificity via traditional immunohistochemistry (IHC) to ensure compatibility and robustness of staining. Working clones were then conjugated as described above, and subject to validation and titration on the MIBI platform as previously described (21). All final images were visually inspected by S.J.R., a board-certified hematopathologist. Fluorophore-conjugated antibodies used for the GeoMx were also validated via immunofluorescence. Details regarding the antibody clones, vendors, conjugated channels, and titers can be found in **Supp Table 10**. Readers of interest are referred to the following publications for a more detailed guide on antibody target selection and optimization (68, 69).

### MIBI Gold slide preparation

The procedure for preparing gold slides was previously described in several studies (67, 70, 71). Briefly, superfrost plus glass slides (Thermo Fisher Scientific, 12-550-15) were first soaked in ddH_2_O and then cleaned by gently rubbing with microfiber cleaning cloths (Care Touch, BD11945) and diluted dish detergent. Subsequently, the slides were rinsed with flowing ddH_2_O to eliminate any residual detergent, then air-dried using a continuous stream of airflow to remove water droplets. The process of coating the slides with 30 nm of Tantalum followed by 100 nm of Gold was performed by the Microfab Shop at Stanford Nano Shared Facility (SNSF) or by New Wave Thin Films (Newark, CA).

### MIBI Slides vectabonding

Coated gold slides were silanized by VECTABOND® Reagent (Vector Laboratories, SP-1800-7) per the protocol provided by the manufacturer. The slides were first quickly rinsed with ddH_2_O to remove dust, then soaked in neat acetone for 5 min, and transferred into 1:50 diluted VECTABOND® reagent in acetone and incubated for 10 min. Slides were then quickly dipped in ddH_2_O multiple times to quench and remove remaining reagents. Remaining water was removed by tapping over Kimwipe without rubbing, air-dried at room temperature (RT) or 37 °C overnight, and stored subsequently at RT.

### MIBI Staining protocol

The procedure of the MIBI staining follows previously described methods (21, 70, 72). Briefly, slides with FFPE sections were baked in an oven (VWR, 10055-006) at 70 °C for 1 hour, then immersed in xylene and incubated for 2 × 10 minutes to thoroughly remove the paraffin. The slides were then subject to a series of solutions for deparaffinization and rehydration using a linear stainer (Leica Biosystems, ST4020): 3× xylene, 3× 100% EtOH, 2×95% EtOH, 1×80% EtOH, 1×70% EtOH, 3× ddH_2_O, 180 seconds each step with constant dipping, and left in ddH_2_O. Antigen retrieval was then performed at 97 °C for 10 minutes with Target Retrieval Solution (Agilent, S236784-2) on a PT Module (Thermo Fisher Scientific, A80400012). After PT Module processing, the cassette containing the slides and solution was allowed to cool to RT on the benchtop. Following a brief minutes rinse with 1× PBS, tissue regions were circled with a PAP pen (Vector Laboratories, H-4000) and then blocked using BBDG (5% normal donkey serum (NDS), 0.05% sodium azide in 1× TBS IHC wash buffer with Tween 20), before an overnight incubation at 4°C with the antibody cocktail (**Supp Table 1**). The next day, slides were subject to 3×5 min washes with the washing buffer (1x TBS IHC wash buffer with Tween 20 and 0.1% BSA) to remove unbound and non-specific antibodies. The samples were briefly rinsed with 1x PBS, fixed with a postfixation buffer (4% PFA + 2% glutaraldehyde in 1×PBS buffer) for 10 minutes, and quenched with 100 mM Tris HCl (pH 7.5). The slides then underwent a series of dehydration steps on the linear stainer (3 × 100 mM Tris pH 7.5, 3× ddH_2_O, 1× 70% EtOH, 1× 80% EtOH, 2× 95% EtOH, 3× 100% EtOH) with 60 seconds for each step. Dried slides were stored in a vacuum desiccator until acquisition.

### MIBI-TOF imaging and image extraction

Datasets were acquired using a commercially available MIBIscope™ system from Ionpath, which is equipped with a Xenon ion source (Hyperion, Oregon Physics). The typical operation parameters for the instrument are listed below:

Production MIBI

Pixel dwell time: 2 ms

Image area: 400×400 µm

Image size: 512×512 pixels

Probe size: ∼400 nm

Primary ion current: Fine mode (5-5.5 nA on a built-in Faraday cup)

Number of planes: 1 depth

The single-channel MIBI images were extracted from raw files generated by the MIBIscope machine using the toffy package (v0.1.0) developed by the Angelo lab https://github.com/angelolab/toffy. For the extraction of most mass channels, a mass range of [-0.25, 0] was employed, while for the 113-Histone H3 channel, a mass range of [-0.25, 0.25] was used.

### MIBI Channel crosstalk removal

Mass-spectrometry based analysis and imaging methodologies such as MIBI can suffer from channel crosstalk caused by adduct formation (71) or isotopic impurities in the elemental labels used. Thus, the Rosetta algorithm was applied to the extracted raw images to remove noise resulting from channel crosstalk with a similar manner as compensating flow-cytometry data https://github.com/angelolab/toffy/blob/main/templates/4a_compensate_image_data.ipynb. Similarly, background signals arising from bare slides or organic fragments can be partially reflected by the gold and “Noodle” background channels. Together, a fine-tuned coefficient matrix was used to remove those channel crosstalk using a local implementation of toffy package (v0.1.0) with minimal modification.

### MIBI Image denoising

The additional noise apart from the channel crosstalks were further removed with a deep learning-based method developed by M.S. and F.M., which poses image denoising as a background-foreground segmentation problem. The underlying concept involves considering the genuine signal as the foreground and the noise as the background. The proposed method employs a supervised deep learning-based segmentation model called UNET (73), to accurately segment the foreground from the given image. To train the model, ground truth data was first generated using a semi-supervised kNN-based clustering method (74). Once the model is trained, it is applied to all markers in all images, producing predicted foreground segmentation maps. These segmentation maps are then multiplied with the original images, eliminating noise and yielding clean images.

### MIBI Cell segmentation

Cell segmentation of the cHL MIBI datasets was performed using a local implementation of deepcell-tf 0.6.0 as described (75, 76). Histone H3 channel was used for the nucleus, while the summation of HLA-DR, HLA1, Na-K-ATPase, CD45RA, CD11c, CD3, CD20, and CD68 was used as the membrane feature. Prior to input into the model, the signals from these channels were capped at the 99.7^*th*^ percentile. The deepcell-tf version used to generate the final segmentation mask, along with the detailed parameters are summarized in **Supp Table 3**.

### MIBI Image intensity normalization

Due to the inherent limitations of the MIBI instrument, an FOV routinely acquired is restricted to a size of 400× 400 µm. Thus, most tiles of the cHL MIBI dataset are composed of multiple stitched adjacent FOVs. Within each tile, the inter-FOV signal level difference and boundary effects were corrected with a series of publicly available scripts as previously described (21, 77).

### MIBI Image to cell expression matrix, REDSEA signal compensation, and across-runs normalization

To generate the cell expression matrix, the counts of each marker within each segmented cell were summed and then divided by the corresponding cell size. This process utilized the normalized stitched TIFs and their respective segmentation masks. To address signal spillover between adjacent cells, REDSEA was applied to the extracted cell expression matrix along with the segmentation mask, as previously described (23). It was observed that Histone H3 signal is positively correlated with most markers, which means that if an FOV has higher Histone H3 signal, the signal of other markers of that FOV would also tend to be higher. Thus, to minimize the unwanted intensity variation between tiles and to make the same channel more comparable across different tiles, the median of Histone H3 counts was found for each tile. Then, the intensity of each marker within each tile was divided by its corresponding median Histone H3.

### MIBI Cell phenotyping

Cell phenotyping of the MIBI dataset was performed through an iterative clustering and annotating process with FlowSOM (78). The scaled cell expression matrix was initially clustered with CD11c, CD14, CD15, CD153, CD16, CD163, CD20, CD3, CD30, CD4, CD56, CD57, CD68, CD8, FoxP3, GATA3, Granzyme B, and Pax-5 to capture most of the cell phenotypes present in the dataset. The resulting clusters were then manually annotated by examining the predominantly enriched markers of each cluster, which was done by plotting Z-score and mean expression heatmaps across all clusters and the phenotypic markers used. Clusters with a clear enrichment pattern were annotated, while clusters with mixed pattern underwent additional rounds of FlowSOM clustering and annotation. To confirm the assigned annotations, Mantis Viewer (79) was utilized. The annotation for each cell cluster was mapped and the raw images of the enriched markers were overlaid for visual inspection. This interactive process was repeated until no additional useful information could be extracted. Cells within clusters that lacked clear enrichment patterns were assigned as “Others”. For the MIBI dataset, a total of 1,538,433 out of 1,669,853 cells (92.2%) were successfully assigned a final annotation. To ensure the accuracy of the annotations, all final annotations were assessed by S.J. and S.J.R..

### Cell neighborhood calling by spatial LDA

Cell neighborhoods were identified using spatial LDA (37), with parameters admm_rho = 0.1, primal_dual_mu = 105, and max_dirichlet_ls_iter = 100. In order to find the optimal number of neighborhoods which strikes a balance between resolution and legibility, we fitted the model with *n* ∈ **N** = {3, 4, 5, 6, 7, 8, 9, 10, 11} neighborhoods and plot-ted a heatmap for cell preference for each model. Starting from *n* = 8, the cell preference of each neighborhood started to stabilize. Meanwhile, a neighborhood with strong preference of “Others” cell type emerged, starting from *n* = 9. Therefore, *n* = 8 was picked as the optimal number of neighborhoods.

### Cell level analysis

#### Expression heatmap and cell type enrichment heatmap

The expression heatmap shows the enrichment of specific markers within a certain cell type when compared with the average expression levels across all cell types. Let **I** = {1, …, *n*} denotes the indices of the lineage and functional markers of interest; **J** = { 1, …, *m*} denotes the indices of the cell types identified; **m**_*i*_ denotes the vector of marker expression for all cells; **m**_*i*,*j*_ denotes the vector of marker expression for all the *j* cell type. The sample mean marker expression for marker *i, µ*_*i*_, the within *j* cell type mean marker expression for marker *i, µ*_*i*,*j*_ , and the sample standard deviation for marker *i, s*_*i*_, were calculated. Then, for each pair of *i* and *j*,

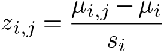

was calculated and presented in the expression heatmap. For the cell type enrichment heatmap shown in **Fig. 3A**, the same procedure was followed except that marker expression was substituted by cell count.

#### Cell phenotype and cell neighborhood enrichment

The cell phenotype enrichment of each cell type *j* was calculated as

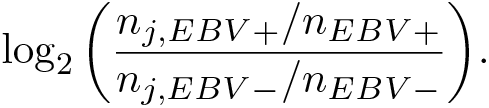

Here, *n*_*j*,*EBV* +_ is the number of cell for cell type *j* among all the EBV+ patients; *n*_*j*,*EBV ™*_ is the number of cell for cell type *j* among all the EBV-patients; *n*_*EBV* +_ and *n*_*EBV ™*_ are the total number of cells among EBV+ and EBV-patients respectively. If a cell type is more prevalent in EBV+ patients than EBV-patients, the log_2_ fold enrichment would be positive. Otherwise, the log_2_ fold enrichment would be negative. If a cell type is equally enriched in both EBV status, the log_2_ fold enrichment would be 0.

#### Tumor score and tumor density classification

For each nontumor cell, a tumor score was calculated based on its distance to tumors within a closed neighborhood of radius *r*. Let **J** = {1, …, *m*} denote the indices of all the tumors in the dataset and *d*_*i*,*j*_ denote the distance from the cell *i* to tumor *j*. Then, the tumor score is calculated as

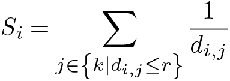

Then, non-tumor cells were classified into tumor dense or tumor sparse based on their tumor score. The cut-off that separated tumor dense and tumor sparse classes was found by identifying the tangent point with the steepest slope to the right of the peak of the distribution of the tumor score.

#### Immune dysfunction level stratified by tumor density and EBV status

Non-tumor cells’ dysfunction level was defined based on combinations of functional markers. For CD4 T cell, CD8 T cell, CD4 CTL, CD8 CTL and Treg, their dysfunction score was defined as

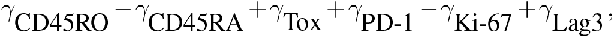

where *γ*marker stands for the intensity of the specified marker. For DC, M1-like, and M2-like cells, their dysfunction score was defined as

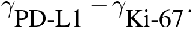

Then, cells were stratified into (tumor dense, *EBV* +), (tumor sparse, *EBV* +), (tumor dense, *EBV*™), and (tumor sparse, *EBV*™). A one-sided paired T test was conducted to compare the dysfunction score between tumor dense and tumor sparse regions within a given EBV status for each of the above mentioned cell types, with the alternative hypothesis that cells in tumor dense regions are more exhausted than those in tumor sparse regions. The test results were corrected for multiple comparisons via Benjamini-Hochberg procedure with a targeted false discovery rate (FDR) of 0.05. The unadjusted p-value and Benjamini-Hochberg procedure statistics and corrected test results can be found in **Supp Table 9**.

### GeoMx Staining protocol

Tissue slides were prepared with modifications from the official Nanostring GeoMx-NGS RNA Manual Slide Preparation protocol. Deparaffinization, rehydration, and antigen retrieval on the two TMA tissue slides were performed using the same procedures as described under the MIBI Staining protocol. Tissue slides were then allowed to cool to RT on the benchtop, and washed in 1× PBS for 5 min at RT. Next, they were digested by Proteinase K (0.5 µg/mL) (Thermo Fisher Scientific, AM2546) for 5 min at 37 °C, and then washed in 1× PBS for 5 min at RT. Subsequently, tissue slides were fixed in 10% neutral buffered formalin (NBF) (EMS Diasum, 15740-04) for 5 min at RT. The fixation process was stopped by incubating twice in 1× NBF stop buffer (0.1 M Tris and 0.1 M Glycine) for 5 min each at RT, followed by a 1× PBS wash for 5 min at RT. Tissue slides were then transferred to fresh 1× PBS while the RNA probe staining cocktail was prepared using the Nanostring RNA Slide Prep kit (Nanostring, 121300313), by combining the Nanostring Human Whole Transcriptome Atlas detection probe (Nanostring, 121401102) set with a custom spike-in panel of probes against 14 targeted EBV genes (*EBER1, EBER2, EBNA1, EBNA2, EBNALP, LMP1, RPMS1, BALF1 BCRF1, BHRF1, BNLF2A, BNLF2B, BNRF1, BZLF1*). The RNA probe staining cocktail was then applied to the tissue slides, sealed with a hybridization cover slip (EMS Diasum, 70329-40), and incubated overnight ( ∼16 hrs) at 37 °C. After RNA probe hybridization, tissue slides were washed twice in Stringent Wash Buffer (2× saline-sodium citrate (SSC) (Millipore Sigma, S6639) in 50% formamide (Millipore Sigma, 344206-1L-M)) for 5 min each at 37°C. Tissues were blocked with Buffer W (Nanostring, 121300313) for 30 min, followed by antibody staining for 1 hr with antibodies against CD3, CD4, and CD45RO, as well as SYTO13 (100 nM) to indicate nucleus and nuclear morphologies. Tissue slides were washed twice in 2× SSC for 2 min each at RT, then stained with anti-mouse antibody for 20 min at RT. Slides were washed twice again in 2× SSC for 2 min each at RT, and then loaded onto the GeoMx to be scanned and selection of region of interest (ROI). Details regarding the antibody clones, vendors, conjugated fluorophores, exposure, and titers can be found in **Supp Table 10**.

### IHC Staining protocol

To guide ROI selection and HRS cell annotation, we performed IHC on an adjacent TMA section for both TMAs with CD30 to identify HRS cells. Deparaffinization, rehydration, antigen retrieval, and cooling to RT for these TMA tissue slides were performed in parallel with those used for GeoMx (see above section). Tissue slides were then washed in UltraPure water (Invitrogen 10977-023) for 5 min at RT, blocked with peroxidase and alkaline phosphatase blocking solution (Vector Laboratories SP-6000-100) for 10 min at RT, and washed with washing buffer (1× TBS-T, 0.1% BSA). Tissue sections were then blocked with 2.5% Normal Horse Serum (Vector Laboratories MP-7500-50) for 1 h at RT, and stained overnight at 4 °C with anti-CD30 antibody diluted in antibody diluent buffer (1× TBS-T, 5% donkey serum, 0.05% sodium azide). Tissue slides were then washed twice in washing buffer, and then stained with secondary anti-mouse IgG and anti-rabbit IgG HRP-conjugated antibodies for 30 min at RT (Vector Laboratories, MP-7500-50). After washing twice in washing buffer for 5 min each, the DAB peroxidase substrate (Vector Laboratories SK-4100) was introduced, and brown coloration was allowed to develop over 2.5 min. Tissue slides were briefly rinsed in tap water and counterstained with hematoxylin (Vector Laboratories H3404-100) for 2 min, and blue coloration was allowed to develop by rinsing in tap water for five times at 3 min each. Finally, tissues were dried in an oven (VWR 10055-006) at 70 °C for 20 min, mounted on a glass coverslip using Vectamount Permanent Mounting Medium (Vector Laboratories H-5000-60), and digitized using a Grundium Ocus®40 microscope (Grundium MGU-00004) to identify CD30-positive stainings. Details regarding the antibody clones, vendors, conjugates, and titers can be found in **Supp Table 10**.

### GeoMx ROI selection

Individual ROIs were selected at regions that contain abundant T cells and HRS cells, by referring to CD3 and CD30 staining patterns respectively, the latter determined through IHC staining on adjacent TMA tissue sections. ROI selections were also optimized to ensure that at least 2 ROIs were obtained from each patient. A total of 126 ROIs (62 EBV-positive, 62 EBV-negative, 2 tonsil) were drawn across both TMAs, and each ROI was drawn as 760×660 µm rectangles (501,600 µm^2^ area each). ROIs were then exported from the GeoMx for cell segmentation and annotation, in order to obtain cell masks that guide transcript extraction from cell populations of interest.

### GeoMx Cell segmentation

Cell segmentation of the GeoMx datasets was performed using a local implementation of deepcell-tf 0.6.0 as described (75, 76). SYTO13 channel was used for the nucleus, while the summation of CD3, CD4, and CD45RO was used as the membrane feature. Prior to input into the model, the signals from these channels were capped at the 99.7^*th*^ percentile. The deepcell-tf version used to generate the final segmentation mask, and the detailed parameters are summarized in **Supp Table 3**.

### GeoMx Cell phenotyping

Regions corresponding to HRS cells were identified based on IHC staining, and HRS cells were manually identified based on nuclear (SYTO13) morphology and adjacent CD30 staining patterns. The visualization of cell nucleus and cells around tumor regions was performed through Mantis Viewer (79). Background signals from CD3, CD4, and CD45RO channels in the GeoMx dataset were first removed by applying percentile cutoffs and visually validated. Cell phenotyping of the GeoMx dataset was performed through PhenoGraph (80), with k = 100 and using CD3, CD4, and CD45RO to identify memory T cells, naive T cells within the TME. The remaining cells which were unidentifiable based on CD3, CD4, and CD45RO were categorized as “other cells”. Finally, Mantis Viewer (79) was utilized to confirm all assigned annotations, where the annotation for each cell phenotype was overlaid over the enriched markers for visual inspection. Binary cell masks corresponding to these 6 cell phenotypes of interest for each ROI were then generated to facilitate downstream transcript extraction through the GeoMx. The marker combinations corresponding to each cell type can be found in **Supp Table 11**.

### GeoMx Transcript extraction and sequencing library preparation

Cell masks were imported into the GeoMx, and transcripts for each cell phenotype across every ROI were aspirated based on the order described in the prior section. In total, 8 collection plates (Nanostring, 100473) were used to collect all aspirates, and aspirates were dried at RT overnight and resuspended in 10 µL of UltraPure water (Invitrogen 10977-023). Each aspirate was then uniquely indexed using the Illumina i5×i7 dual indexing system through Nanostring NGS library preparation kits (Nanostring, 121400201, 121400202, 121400203, 121400204). The PCR reaction was prepared in 96-well plates, where each well contained 4 µL of aspirate, 1 µM of i5 primer, 1 µM of i7 primer, and 1x library preparation PCR Master Mix, adding up to 10 µL in total volume. The PCR reaction conditions were 37 °C for 30 min, 50 °C for 10 min, 95 °C for 3 min, followed by 18 cycles of 95 °C for 15 s, 65 °C for 60 s, 68 °C for 30 s, followed by a final extension of 68 °C for 5 min before holding at 4 °C. Next, 4 µL of PCR products from each plate were then pooled into DNA LoBind tubes (Eppendorf 022431021) for purification, where 1.2× volume of AMPure XP beads (Beckman Coulter A63881) were first added to the pooled PCR products and at RT for 5 min. Beads were then pelleted on a magnetic stand (Thermo Fisher Scientific 12321D), washed twice with 1 mL of 80% ethanol, and eluted with 54 µL of elution buffer (10 mM pH 8.0 Tris-HCl, 0.05% Tween-20). Another round of purification was performed using 50 µL of eluted DNA in the same approach, using 1.2× volume of AMPure XP beads and washing twice in 1 mL of 80% ethanol. A final elution was done at 2:1 ratio of aspirate (number of wells) to elution buffer (volume in µL), and 0.2 µL of the final eluate was diluted in 9.8 µL of UltraPure water (Invitrogen 10977-023) (1:10 dilution) to confirm library purity through Agilent BioAnalyzer. Finally, libraries were paired-end sequenced on NovaSeq6000 with a sequencing depth of ∼7.5 billion reads, which is at least 1.2× greater than recommended by the official protocol.

### GeoMx Data processing

#### Transcript mapping and counting

The NGS barcodes from the Nanostring human WTA panel and custom EBV-specific probes were mapped and counted using the commercial GeoMx Data Analysis software pipeline (19), using FASTQ files generated from NGS sequencing. Subsequently, the ‘.dcc’ files produced by the GeoMx Data Analysis software pipeline were used as input to generate the gene-level counts table. R package GeomxTools (v.3.6.2) with default setting was implemented, and raw gene counts table was produced and used for normalization and batch effect correction.

#### Data normalization and batch effect removal

The standR (v.1.4.2) workflow included normalization, negative control gene (NCG) searching, and batch effect removal using the RUV4 method (81), and thus was used to normalize raw gene counts and reduce patient-level batch effects induced by individual variability (81). To fit the best practice of batch effect removal, a grid search was implemented to find the optimal hyperparameter combinations that minimized individual variations while keeping EBV condition and cell type variations. By following the standR workflow, four normalization methods were adopted, including the trimmed mean of M-values (TMM), log counts-per-million reads (CPM), quantile normalization, and size factor normalization. Nine grids of the number of NCG genes were selected (100, 300, 500, 700, 900, 1100, 1300, and 1500). The five grids of the number of k-covariance matrices for the RUV4 method were set to 3, 5, 7, 9, and 11. In total, 180 parameter sets were evaluated.

#### Assessing the data after batch effect removal

The effect of batch correction was assessed using the silhouette and the kBET scores (82), as implemented in the R package kBET (version 0.99.6). The silhouette score quantifies cluster separation, with a higher score signifying more distinct clustering. The kBET assessment involved quantifying the rejection rate through Pearson’s chi-square test and comparing the distribution of local and global batch labels among the k-nearest neighbors. Specifically, a high kBET score of a factor indicated that less bias was introduced by this factor. Both silhouette and kBET scores were used to evaluate the consistency across all samples in terms of patient factor for individual variations, cell type, and EBV condition factor for biological variations. Therefore, the objective of the standR workflow’s grid search was to identify a parameter set that minimized the batch effect related to the patient factor while maximizing the differentiation by cell type and EBV condition in the following steps. First, post-corrected expression matrices were derived based on the standR pipeline under a specific parameter set. Second, silhouette and kBET scores for each parameter set were computed according to three factors. Third, adopting the concept from the previous study (83), silhouette and kBET scores were ranked in following rules. Specifically, scores determined by the patient factor were ranked in ascending order, where the lowest score received ranked one, and the highest score was assigned ranked 180. Scores determined by cell type and EBV condition were ranked in decreasing order, where the highest score received ranked one, and the lowest score was assigned ranked 180. Lastly, an overall rank was calculated by averaging the rank of three factors and two scores, and the optimized parameters were selected based on the overall rank. The batch effect corrected data was then used for all subsequent analysis. The detailed evaluation scores during the batch effect correction process can be found in **Supp Table 14**, which provides an assessment of different parameters set against various factors (e.g. patient, cell type (CT), and EBV status), while incorporating both kBET and silhouette scores as well as their corresponding ranks to evaluate batch effect correction.

### GeoMx Data analysis

#### General analysis related to gene expression

To identify significantly upregulated genes in each of the 6 region types (CD4 memory, CD8 memory, CD4 naive, CD8 naive, Tumor, and Other), the function ‘findallmarkers’ from the R package ‘seurat’ were implemented. For visualization, only the top 30 significant genes (*p*.*adj* < 0.05) with the highest log-fold change were plotted in the heatmap. Differentially expressed genes were identified by R package ‘limma’ with default parameters. G2M scores were calculated in the Tumor regions, with function ‘CellCycleScoring’ from R package ‘seurat’. GSVA scores (gene pathway scores) were calculated using the function ‘gsva’ from R package ‘GSVA’ (v.1.42.0) with default parameters.

#### Receptor-ligand interaction analysis

Statistical inference of chemokine receptor-ligand interactions between tumor and T cell regions were performed via CellphoneDB v5 (44, 45) using the repository of chemokines curated within. Briefly, after quantifying the co-expression of receptor-ligand pairs between T cell and tumor regions within each ROI, an expression heatmap was generated based on the mean coexpression score across all ROIs. Statistical significance was assessed by comparing receptor-ligand interactions against a null distribution that was generated by simulating random receptor-ligand interactions across 1000 iterations.

#### Circos plots

To determine the association between the targeted pathway and its corresponding genes, this analysis involved the assessment of each gene’s contribution to a specified biological pathway using Spearman’s rank correlation coefficient. The samples were first stratified based on EBV condition and cell types, and then Spearman correlation coefficients were computed for these subgroups. A Circos plot, implemented by the circlize R package (version 0.4.15), was subsequently utilized to graphically represent the relationships between gene expression and pathway activity.

### Data visualization

Single channel and multi-color images were assembled with ImageJ. Visualizations of the analysis results were either produced using Excel, or R packages ‘ggplot2’ and ‘complexheatmap’.

### Code availability

All code for analysis and data visualization can be downloaded at https://github.com/SizunJiangLab/Hodgkin_EBV_MIBI.

## ACKNOWLEDGEMENTS

We thank Matthew Newgren and Maciej Zerkowski previously from Ionpath for their unwavering technical support, members of the Shipp, Rodig, and Jiang labs for helpful discussions, and Marvin Nayan, Adam Limb, and Mike Chen from Nanostring for GeoMx technical support.

S.J. is supported by NIH DP2AI171139, P01AI177687, R01AI149672, a Gilead’s Research Scholars Program in Hematologic Malignancies, a Sanofi Award, the Bill & Melinda Gates Foundation INV-002704, the Dye Family Foundation, and previously by the Leukemia Lymphoma Society Career Development Program. G.P.N. is supported by the Rachford and Carlota A. Harris Endowed Professorship. F.M. is funded by R35GM138216 and the Fredrick National Laboratory. B.E.G. is funded by NIH CA275301 and CA228700. S.J.R. is supported by Bristol-Myers Squibb (BMS) International Immuno-Oncology Network (II-ON). S.J.R. and M.A.S. are supported by a Blood Cancer Discoveries Grant Program from the Leukemia Lymphoma Society, The Mark Foundation, and The Paul G. Allen Frontiers Group. This article reflects the views of the authors and should not be construed as representing the views or policies of the institutions that provided funding.

## AUTHOR CONTRIBUTIONS

S.J., S.J.R., M.A.S. conceived of the study and planned the experiments with input from all authors. Y.Y.Y., Y.B., H.A.M., K.W., H.C., S.J. performed experiments. H.Q., Y.B., Y.C., B.Z., Y.Y.Y., J.Y., S.J. performed data analysis. K.W., M,S., S.S., D.N., V.S., P.R., S.P.T.Y., P.C., J.P., P.S., G.L., A.Y.H., H.W., H.C., L.F., B.M., B.E.G., C.M.S., B.Z., G.P.N., B.Z., A.K.S., M.A., F.M., Q.M., W.R.B. contributed to the samples, reagents, or pathological and/or technical expertise. Work performed by G.L. was conducted during his tenure as a visiting member of the Jiang laboratory. V.S., K.W., P.R., W.R.B., S.J.R. performed the pathological analysis for the archival human cohort used in this paper. Y.Y.Y., H.Q., Y.B., B.Z, Y.C., M.A.S., S.J.R., S.J. wrote the manuscript with input from all authors. S.J., S.J.R., M.A.S., provided supervision and funding acquisition. All authors reviewed and edited the final manuscript.

