## Supplementary Figures for "Epstein-Barr Virus Orchestrates Spatial Reorganization and Immunomodulation within the Classic Hodgkin Lymphoma Tumor Microenvironment"

[1]Center for Virology and Vaccine Research, Beth Israel Deaconess Medical Center, Harvard Medical School, Boston, MA, United States
[2]Program in Virology, Division of Medical Sciences, Harvard Medical School, Boston, MA, United States
[3]Department of Pathology, Stanford University, Stanford, CA, United States
[4]Department of Chemistry, Stanford University, Stanford, CA, United States
[5]Department of Microbiology and Immunology, Stanford University, Stanford, CA, United States
[6]Pelotonia Institute for Immuno-Oncology, The Ohio State University, Columbus, Ohio, United States
[7]Department of Pathology, Brigham and Women's Hospital, Harvard Medical School, Boston, MA, United States
[8]Broad Institute of Harvard and MIT, Cambridge, MA, United States
[9]Department of Pathology and Laboratory Medicine, University of Rochester Medical Center, Rochester, NY, USA
[10]Department of Medical Oncology, Dana-Farber Cancer Institute, Harvard Medical School, Boston, MA, United States
[11]Université de Paris, Assistance Publique-Hôpitaux de Paris, Hemato-oncologie, Saint-Louis Hôspital, Paris, France
[12]Center of Hepato-Pancreato-Biliary Surgery, The First Affiliated Hospital of Sun Yat-sen University, Guangzhou, Guangdong, People's Republic of China
[13]Division of Infectious Disease, Department of Medicine, Brigham and Women's Hospital and Harvard Medical School, Boston, MA, United States
[14]Department of Pathology and Neuropathology, University Hospital and Comprehensive Cancer Center Tübingen, Tübingen, Germany
[15]Cluster of Excellence iFIT (EXC 2180) "Image-Guided and Functionally Instructed Tumor Therapies", University of Tübingen, Germany
[16]Department of Chemistry, Massachusetts Institute of Technology, Cambridge, MA, United States
[17]Department of Biomedical Informatics, The Ohio State University College of Medicine, Columbus, Ohio, United States
[18]Department of Pathology, Dana Farber Cancer Institute, Boston, MA, United States
[*]Equal Contribution
[†]Senior Authors

### Supplementary Figures

#### MIBI staining: Antibody Validation

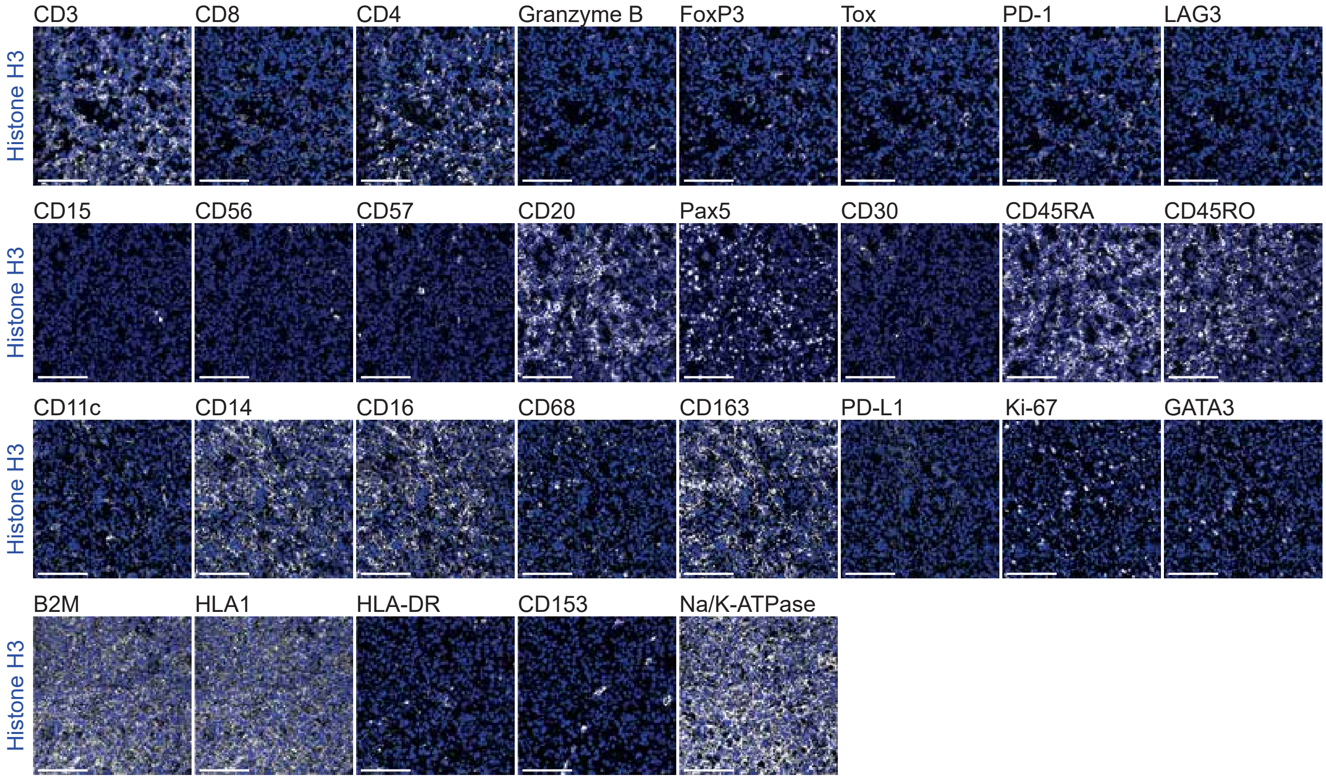


**Figure S1, related to Figure 1. Validation of MIBI antibody staining specificity.** Representative MIBI images across cHL tissue sections, showing 29 antibody markers (white) overlaid with the cell nucleus antibody marker Histone H3 (blue). Scale bar: 100 µm.

### Phenotype Maps



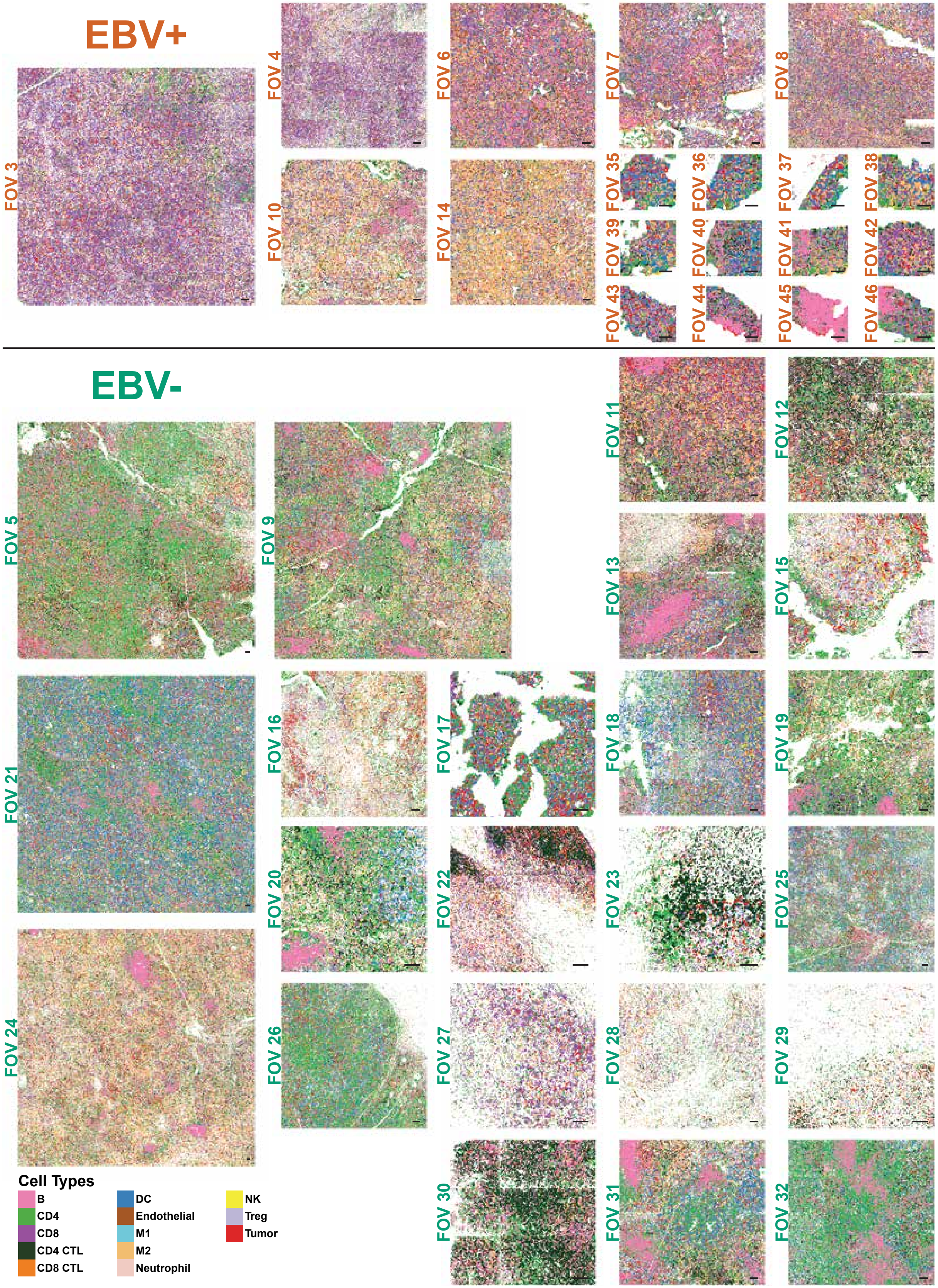


**Figure S2, related to Figure 2. Validation of cell phenotyping from MIBI images.** Phenotype maps of all the MIBI-acquired FOVs across cHL tissue sections, generated through iterative clustering and annotation based on single-cell MIBI features. Scale bar: 100 µm.

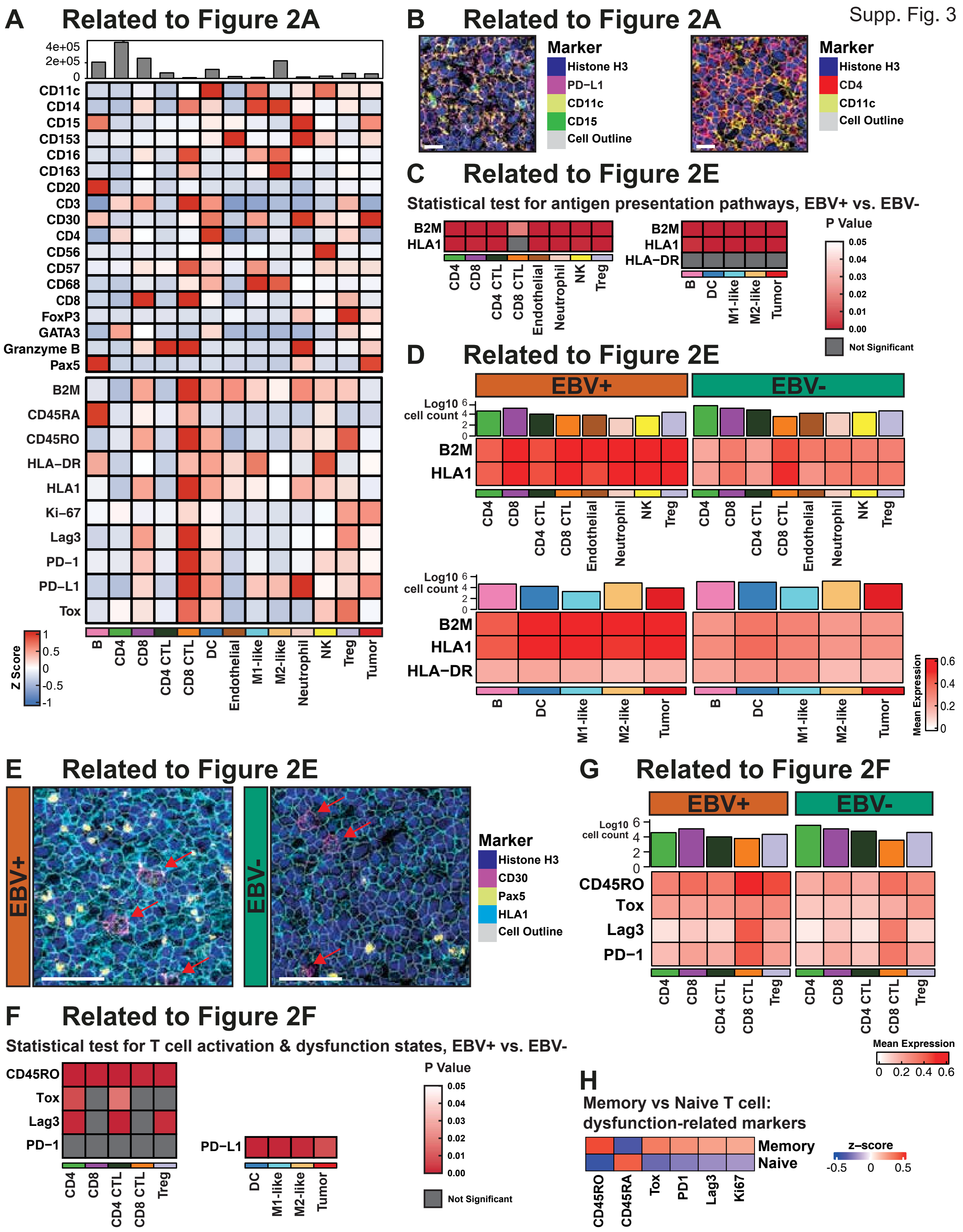


**Figure S3, related to Figure 2. Validation of marker expression on a cell type level. (A)** Relative z-score expression levels of phenotypic (top) and functional markers (bottom) for the annotated cell phenotypes in the EBV-positive and EBV-negative cHL MIBI dataset. **(B)** Representative MIBI images showing neutrophils (CD15) close to immune constituents that have express PD-L1, such as dendritic cells (CD11c) (left), and dendritic cells (CD11c) in close contact with CD4 T cells (CD4) (right). Cell outlines are shown to indicate segmented cell boundaries. Scale bar: 20 µm. **(C)** and **(F)** Test results for **Figs. 2E** & **2F**, respectively. P values were generated from one-sided Wilcoxon rank sum tests, with the alternative hypothesis that the distribution of a given marker for the EBV+ population is shifted to the right of the distribution for the EBV- population. The test results were adjusted for multiple comparisons using the Benjamini-Hochberg method with a targeted FDR at 0.05. Unadjusted p-value and Benjamini-Hochberg corrected test results are in **Supp Table 7**. **(D)** and **(G)** Relative mean expression level heatmaps, a re-representation of relative z-score expression level heatmaps in **Figs. 2E** & **2F**. **(E)** Representative MIBI images showing MHC Class I (HLA1) expression differences between the EBV-positive and EBV-negative cHL TME, with additional markers for B cells ($Pax5^{hi}$) and HRS cells (CD30, $Pax5^{lo}$) shown. Red arrows point to HRS cells. Cell outlines are shown to indicate segmented cell boundaries. Scale bar: 50 µm. **(G)** Expression heatmap of immune exhaustion markers on T cells stratified into memory and naive populations based on the relative expression of CD45RO and CD45RA. The data used to generate this heatmap is in **Supp Table 15**.

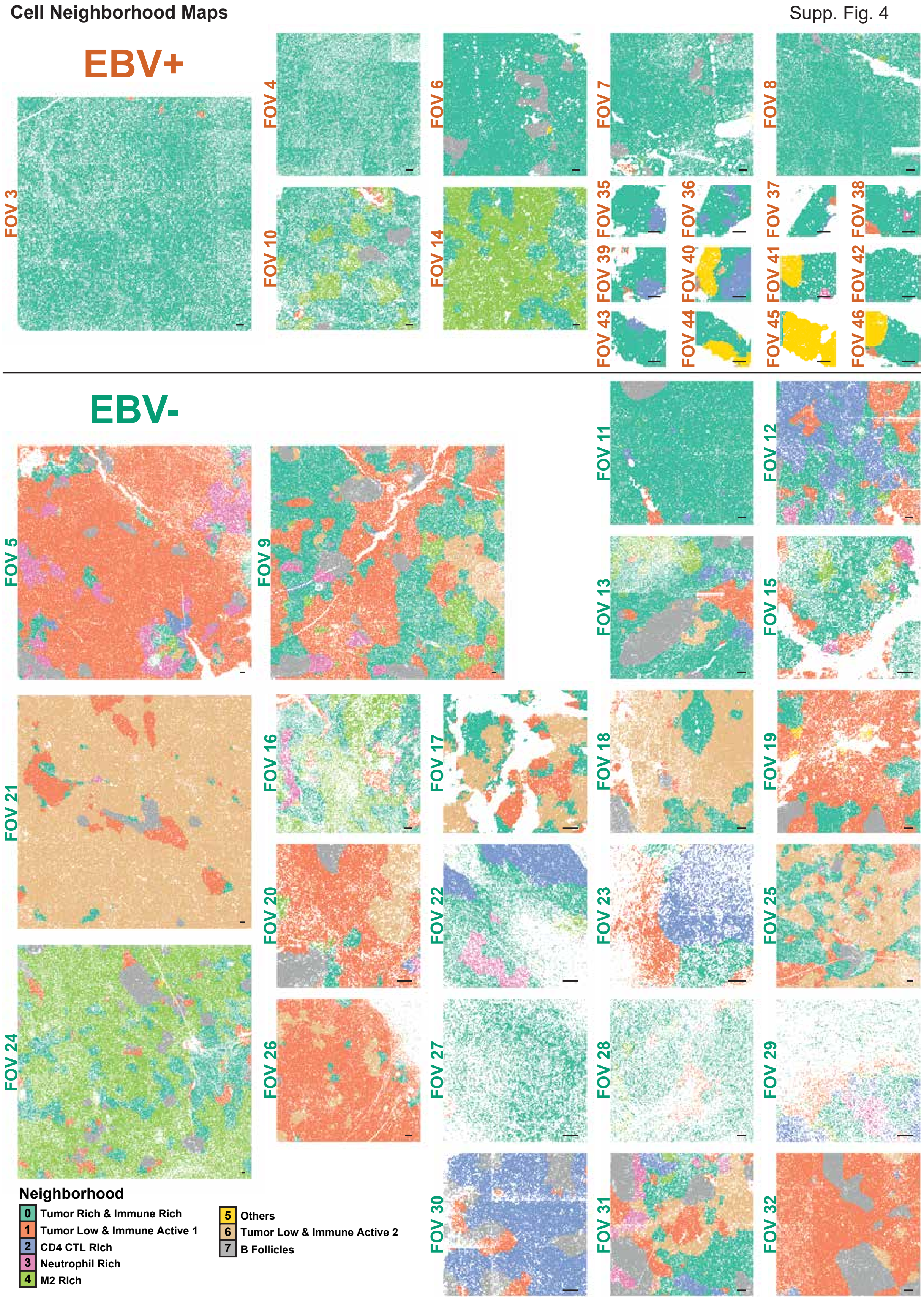


**Figure S4, related to Figure 3. Validation of cell neighborhood identification from annotated cell types.** Cell neighborhood maps of all MIBI-acquired FOVs across cHL tissue sections, generated through spatial LDA (1) based on annotated cell phenotypes (see **Supp Fig. S2**). Scale bar: 100 µm.

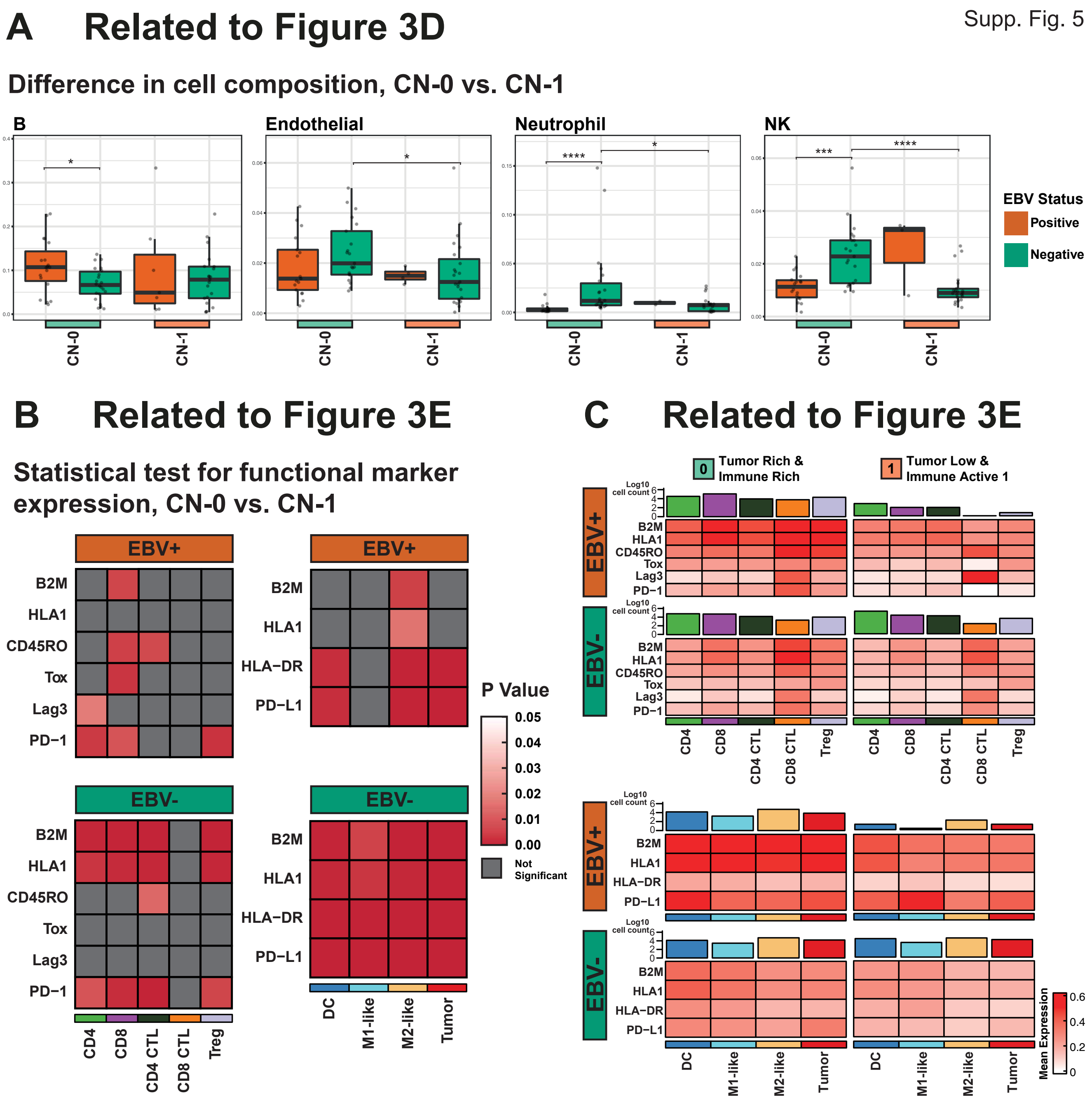


**Figure S5, related to Figure 3. Statistical test results for comparison of cell proportions and functional marker expression on a cell neighborhood level. (A)** Relative proportion of other cell types within CN-0 and CN-1, stratified by EBV status. Significance stars are only shown for statistically significant comparisons ($p \leq 0.05$). **(B)** Test results for **Fig. 3E**. P values were generated from two-sided Wilcoxon rank sum tests, with the alternative hypothesis that within the EBV+ or EBV- population respectively, the distribution of a given marker is different across CN-0 and CN-1. The test results were adjusted for multiple comparisons using the Benjamini-Hochberg method with a targeted FDR at 0.05. Unadjusted p-value and Benjamini-Hochberg corrected test results are in **Supp Table 8** and **9**. **(C)** Relative mean expression level heatmap, a re-representation of relative z-score expression level heatmap in **Fig. 3E**.

### Tumor Score Maps

Supp. Fig. 6

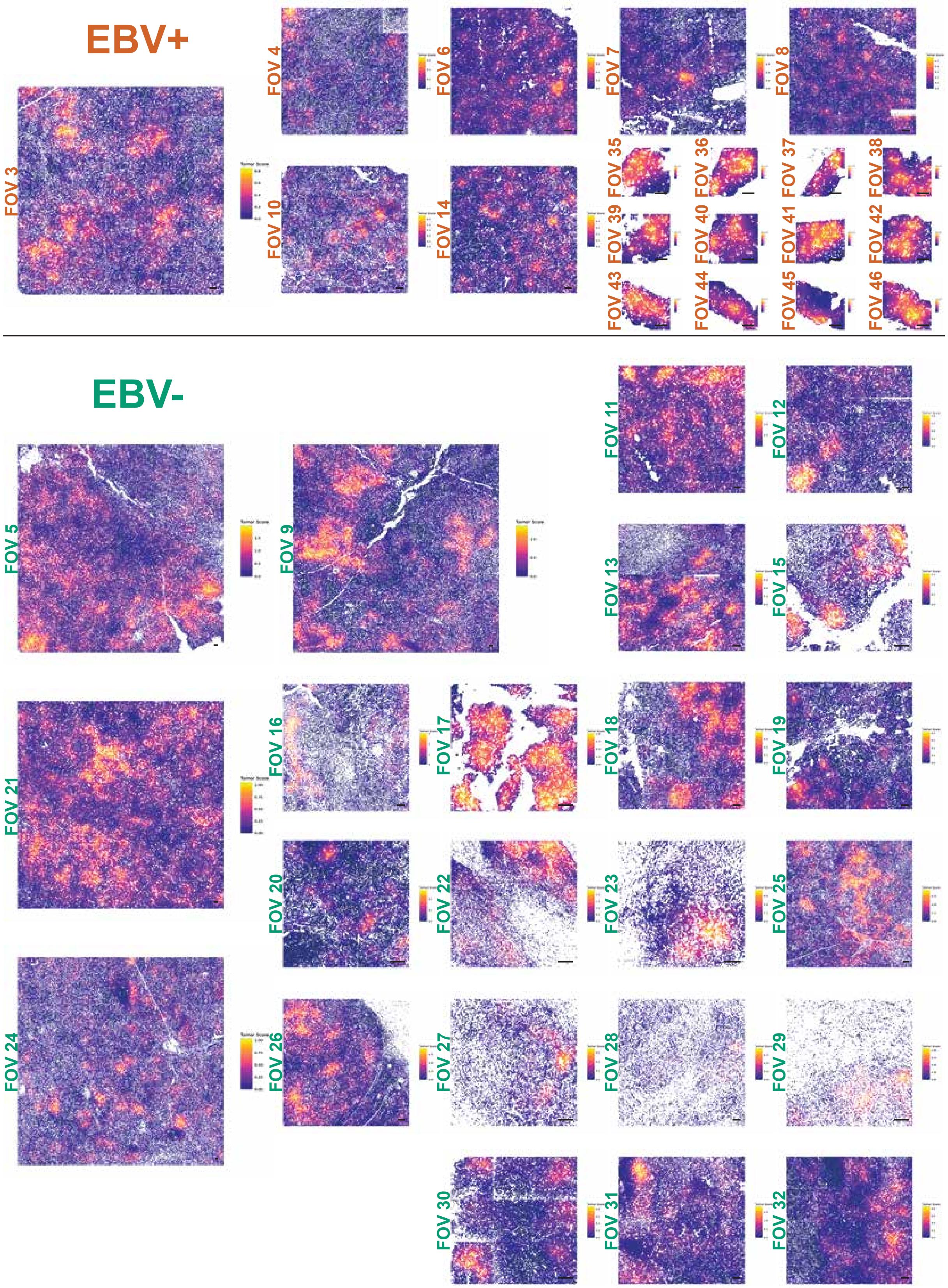


**Figure S6, related to Figure 4. Validation of tumor score metric.** Tumor score maps of all MIBI-acquired FOVs across cHL tissue sections, generated based on the spatial proximity to tumor cells (see **Materials & Methods**). HRS cells are not considered in the tumor score metric and do not have a color assigned. Scale bar: 100 µm.

Tumor Dense / Sparse Map

Supp. Fig. 7

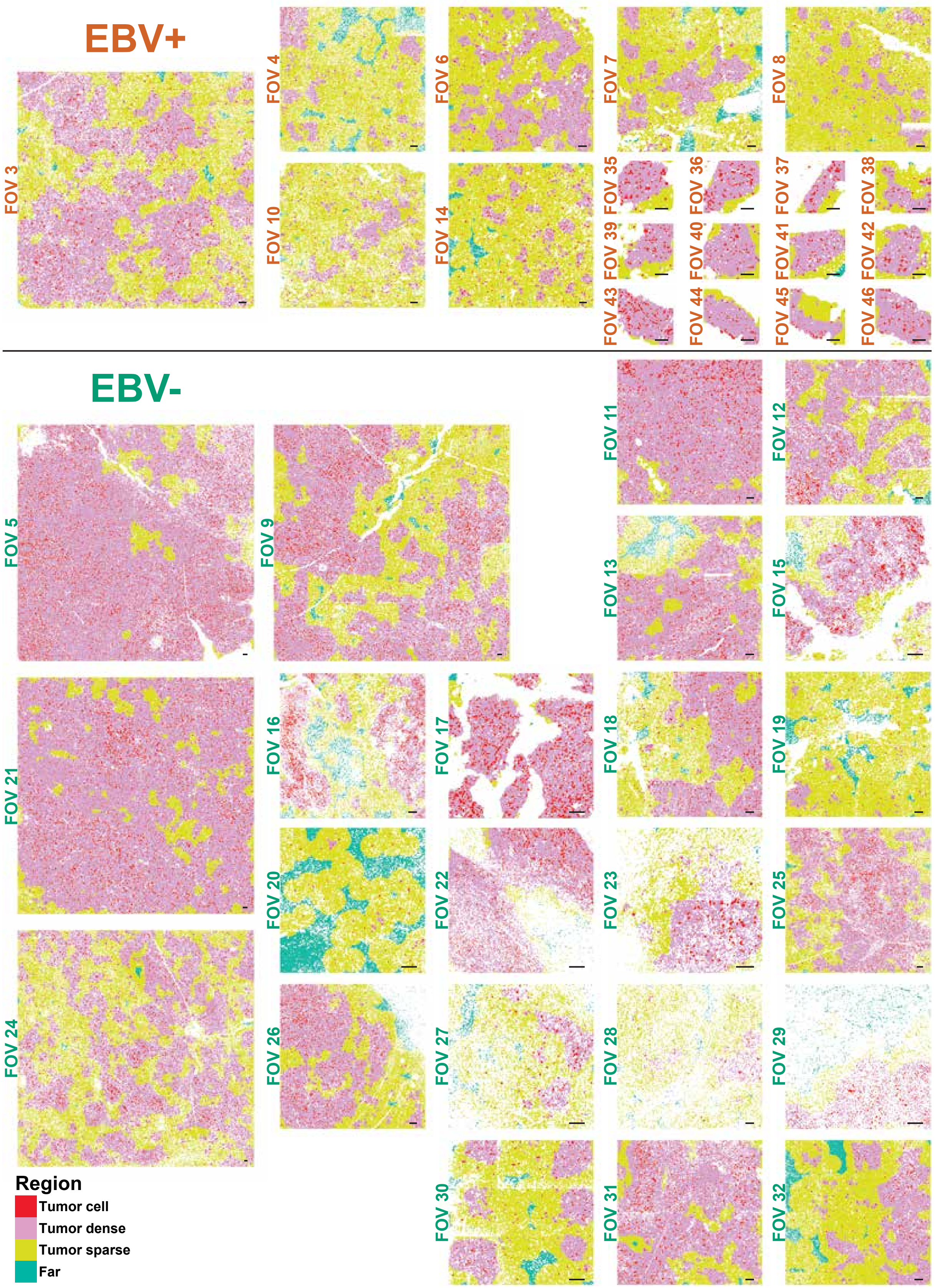


**Figure S7, related to Figure 4. Validation of tumor dense/sparse region stratification.** Tumor dense and tumor sparse maps of all 34 MIBI-acquired FOVs across cHL tissue sections, generated based on the tumor score metric of each non-tumor cell (see **Materials & Methods**). Scale bar: 100 μm.

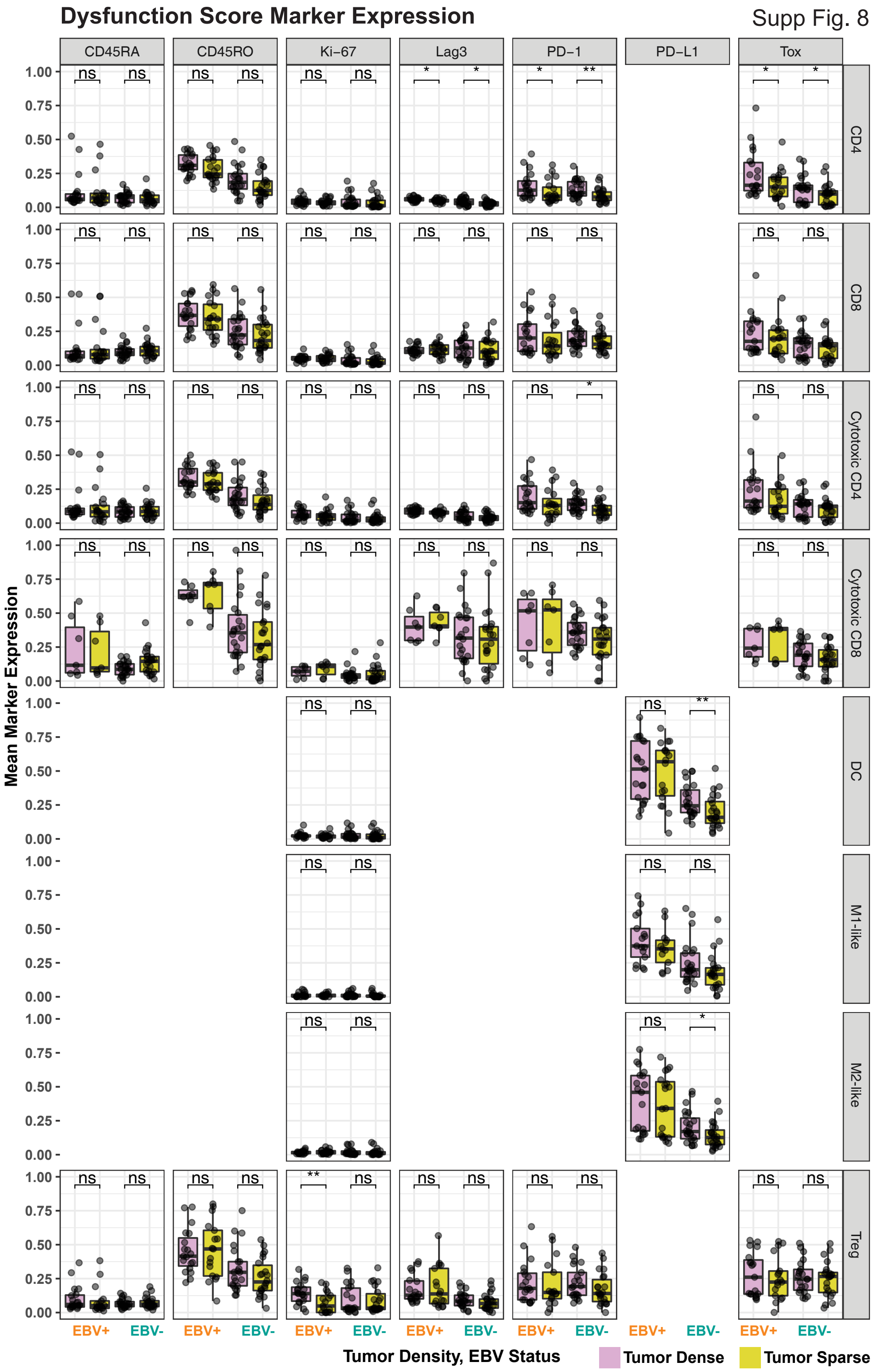


**Figure S8, related to Figure 4. Statistical test results for comparison of functional marker expression on a tumor dense/sparse level.** Relative expression of markers that comprise the exhaustion score metric. For each tumor dense and tumor sparse pair, a one-sided T test was performed, with the alternative hypothesis that the difference between the mean exhaustion score of the tumor dense region in an FOV and that of the tumor sparse region is greater than 0 (* $p \leq 0.05$, ** $p \leq 0.01$, *** $p \leq 0.001$, **** $p \leq 0.0001$). The test results were adjusted for multiple comparisons using the Benjamini-Hochberg method with a targeted false discovery rate (FDR) at 0.05. Unadjusted p-value and Benjamini-Hochberg corrected test results are in **Supp Table 10**.

Supp Fig. 9

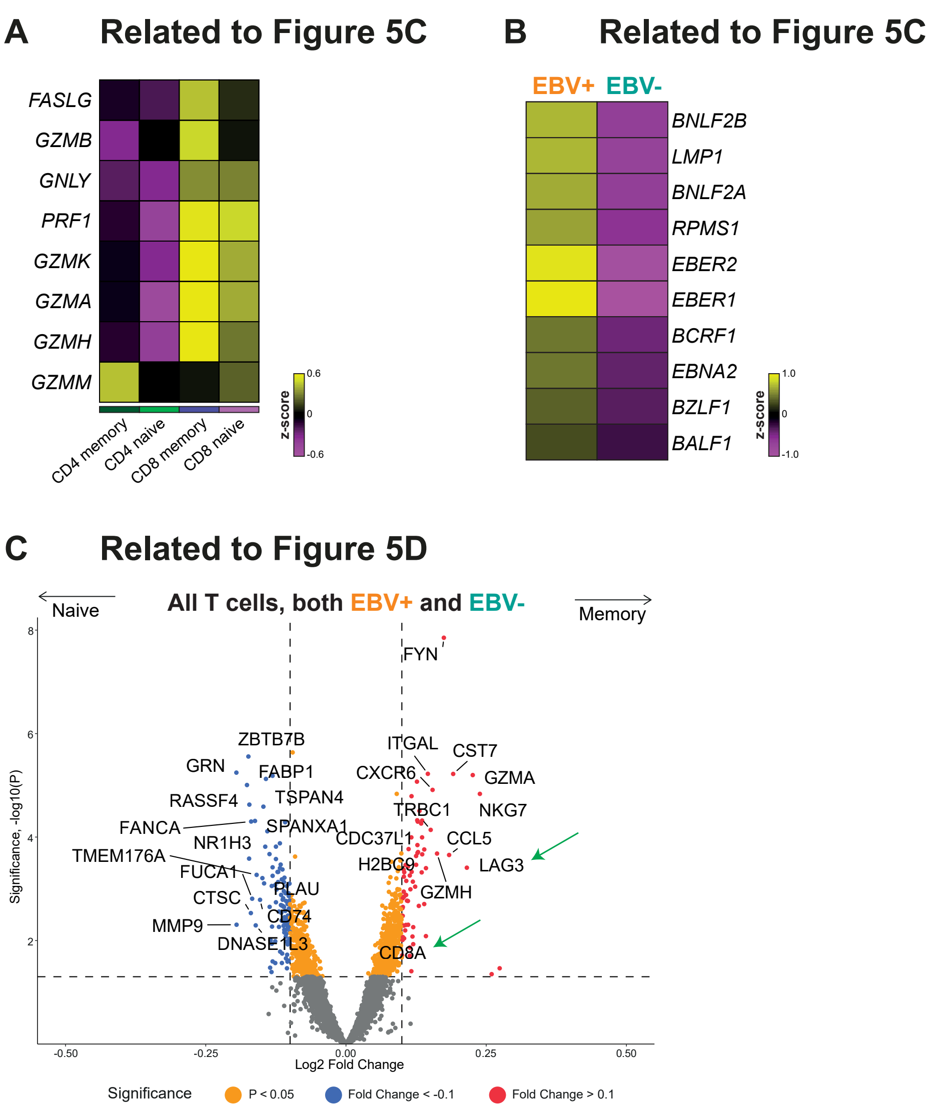


**Figure S9, related to Figure 5. Comparison EBV-linked transcriptional differences. (A)** Expression heatmap of cytotoxic transcripts associated with each annotated cell region. **(B)** Expression heatmap of EBV genes in EBV-positive and EBV-negative tumor regions. **(C)** Volcano plot of memory (CD45RO+) vs. naive (CD45RO-) T cells, with some of the most differentially expressed genes shown. *CD8A* and *LAG3* are indicated by the green arrows.

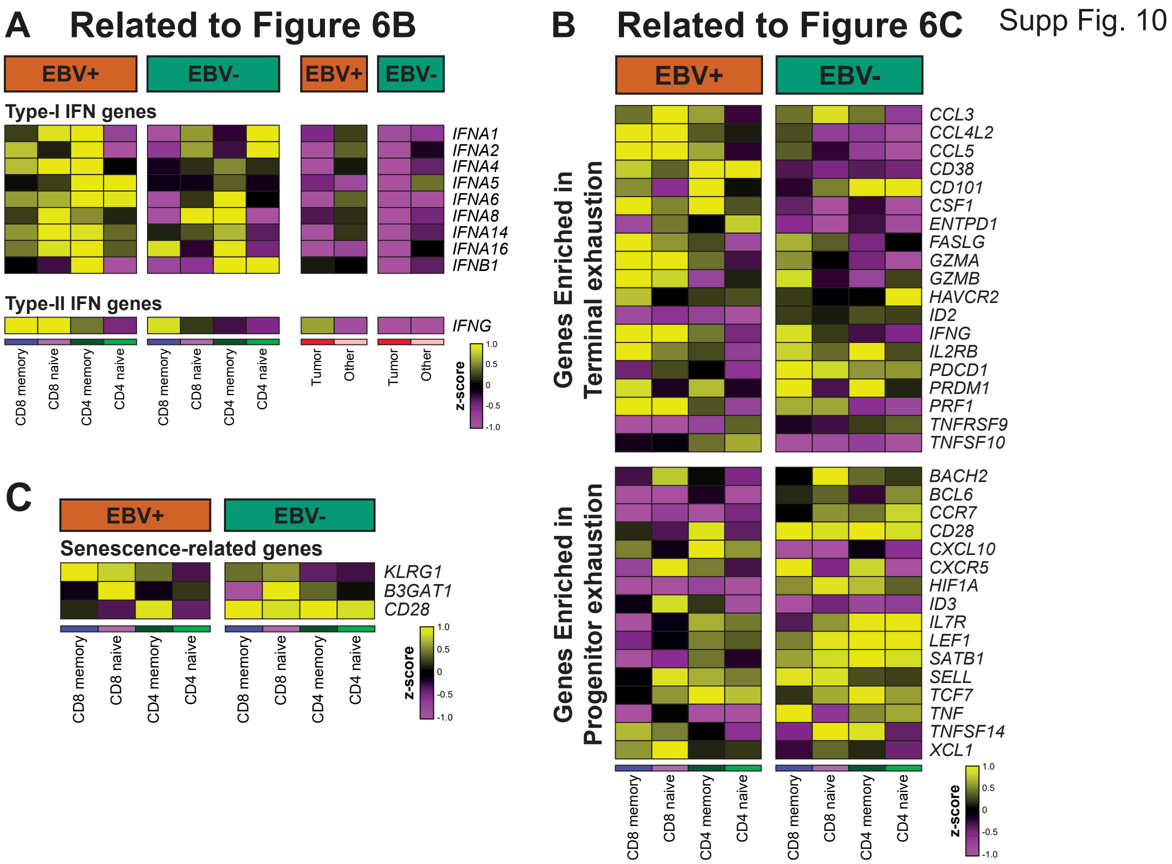


**Figure S10, related to Figure 6. Expressions of genes used to score specific gene sets. (A)** Expression heatmap of interferons used to score “Type-I IFN genes” and “Type-II IFN genes” pathways in **Fig. 6B**. This list does not include all known interferons because it only contains probes available in the GeoMx probe set. **(B)** Expression heatmap of key differentially expressed genes between terminal and progenitor T-cell exhaustion states (see Fig. 2D in (2)). **(C)** Expression heatmap of genes and pathways associated with T-cell senescence (3).

### Related to Figure 6D

Supp Fig. 11

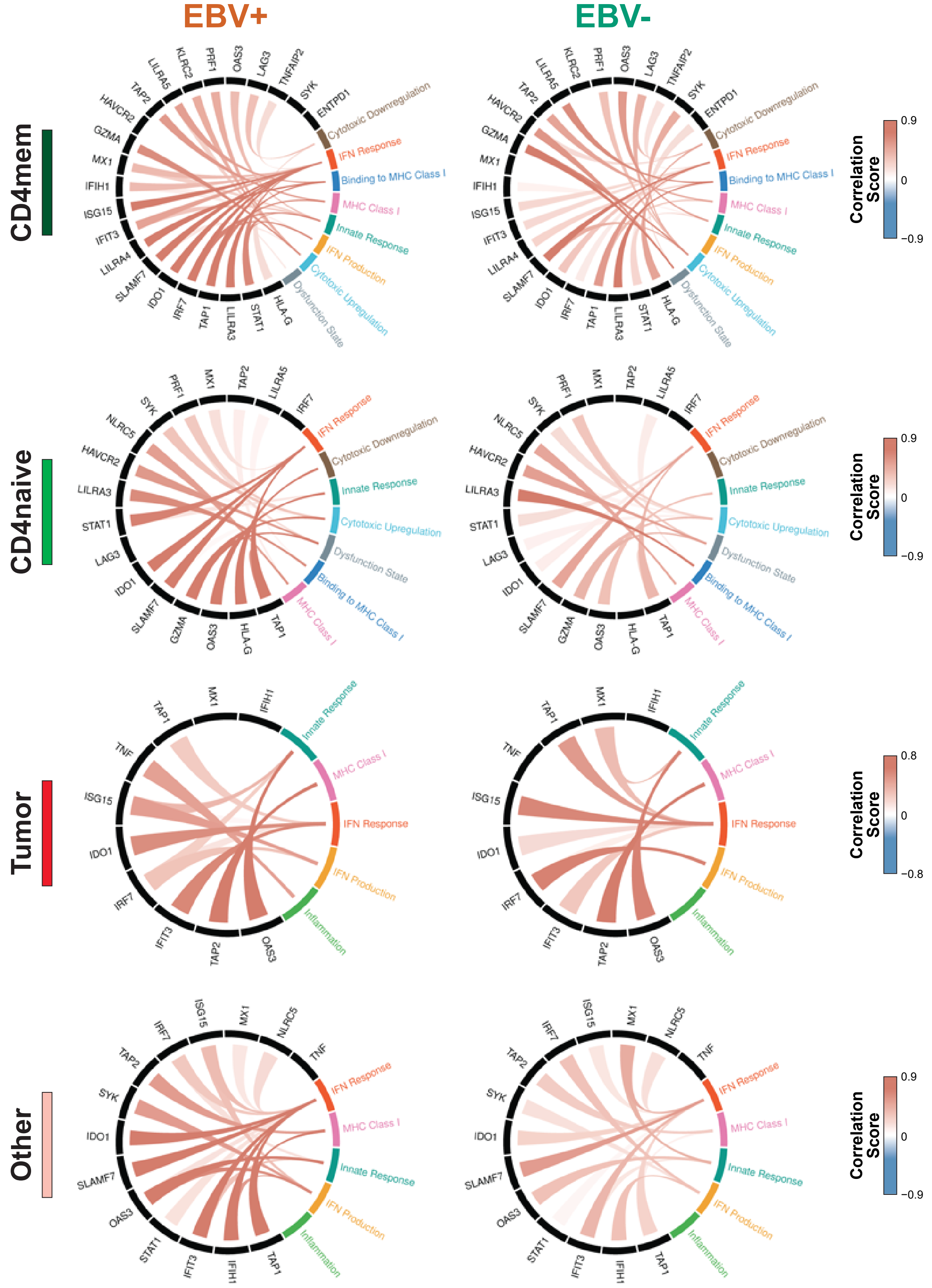


**Figure S11, related to Figure 6. Comparison of gene-pathway correlations for all other cell types, stratified by EBV status.** Circos plot for all other cell types, in addition to those shown in **Fig. 6D**.
